# Combining phenomic and genomic selection for pea breeding improvement

**DOI:** 10.1101/2025.08.06.668650

**Authors:** Anthony Klein, Virgilio Freitas, Abdou Wade, Grégoire Aubert, Myriam Naudet-Huart, Michael Touratier, Jennifer Régulier, Jean-François Herbommez, Catherine Kraut, Guillaume Roullet, Aude Carlier-Lézie, Nadim Tayeh, Renaud Rincent, Philippe Dufour, Judith Burstin, Gaëtan Touzy

## Abstract

Pea (*Pisum sativum* L.) is a strategic crop in the development of sustainable agriculture. However, the genetic gain remains limited despite advances in breeding. Genomic selection holds promise to accelerate varietal improvement, but its high implementation cost restricts its use in crops. Phenomic selection, based on near-infrared spectroscopy data, is a cost-effective alternative demonstrated in various crops, but not yet undertaken in pea. This study aims to assess the predictive ability of phenomic selection, alone and combined with genomic selection, for yield-related traits in a panel of elite spring pea lines evaluated across twelve environments. Three cross-validation scenarios were implemented to simulate predictions across different years and locations. Our results show that phenomic prediction is as effective as genomic selection at predicting yield, and is more accurate for seed protein content. The integrative model, combining spectral and molecular data, consistently achieved the highest accuracy for most traits, particularly for complex traits such as grain yield and seed protein. In temporal prediction scenarios, the most accurate predictions were obtained using the spectra data from the same year as phenotyping. In spatial prediction scenarios, predictive accuracy varied by site and year, nevertheless, integrative phenomic-genomic models consistently outperformed univariate approaches. These findings confirm the potential of phenomic selection in pea and underscore the added value of combining near-infrared spectroscopy and genotyping data to improve the prediction of complex traits in breeding programs. In the face of increasing environmental variability, the integrative approach offers a valuable tool for accelerating genetic gain.

**Key message:** The integration of spectral data into prediction models enhances the predictive ability for complex traits in pea.

## Introduction

Pea (*Pisum sativum* L.) is a cool-season pulse crop and a key component for sustainable cropping systems (Champ et al. 2015). Pea seeds are an important source of proteins and provide an exceptionally varied nutrient profile (Burstin et al. 2011). Beyond its agronomic value, pea research has led to major advances in plant genetics (Singh et al. 2023; Smýkal et al. 2016) and been boosted by the development of genetic resources (Alves-Carvalho et al. 2015; Duarte et al. 2014; Kreplak et al. 2019; Tayeh et al. 2015a). Numerous studies have explored the genetic determinism of complex traits in pea, including seed yield and quality (Bourgeois et al. 2011; Gali et al. 2024; Gali et al. 2019; Klein et al. 2020; Tar’an et al. 2004), resistance to biotic stresses (Aznar-Fernández et al. 2020; Barilli et al. 2020; Coyne et al. 2019; Desgroux et al. 2016; Jain et al. 2015; Jha et al. 2017; Lavaud et al. 2024; Leprévost et al. 2023) and tolerance to abiotic stresses (Beji et al. 2020; Huang et al. 2023; Iglesias-García et al. 2015; Klein et al. 2014; Lejeune-Henaut et al. 2008; Tafesse et al. 2021). Recently, the availability of high-quality reference genomes for multiple Pisum accessions has provided new resources to improve pea breeding (Kreplak et al. 2025; Liu et al. 2024; Yang et al. 2022).

Despite advances in research and in breeding, genetic progress in pea breeding remains limited, estimated at 0.067 t/ha/year for the French spring pea and 0.054 t/ha/year for the French winter pea (van Boxsom and Retailleau 2022), likely due in part to significant genotype-by-environment interactions and to limited research investment in this minor crop (Fugeray-Scarbel and Lemarié 2024). Intensified breeding efforts represent a key strategy to enhance the profitability of pea cultivation and promote its large-scale development (Magrini et al. 2016). In this context, in addition to marker-assisted selection, the development of genetic and genomic tools has enabled the emergence of Genomic Selection (GS) programs in pea (Tayeh et al. 2015b). Key targeted agronomic traits, included grain yield and thousand-seed weight (Annicchiarico et al. 2019; Burstin et al. 2015; Crosta et al 2025; Crosta et al. 2023; Tayeh et al. 2015c), as well as tolerance to biotic and abiotic stresses (Carpenter et al. 2018; Gondalia et al. 2022; Osuna-Caballero et al. 2024; Parihar et al. 2022). GS is proposed as a powerful approach to increase genetic gain (Bernardo and Yu 2007; Heslot et al. 2015; McGaugh et al. 2021; Meuwissen et al. 2001; Xu et al. 2020), particularly by reducing the breeding cycle length in annual inbred crops. By predicting Genomic Estimated Breeding Values (GEBV), GS enables earlier and more accurate selection of superior genotypes (Li et al. 2022). However, the efficiency of the GS in plant breeding varies significantly, depending on the trait genetic architecture, specific breeding and cultivation systems (Akdemir and Isidro-Sánchez 2019; Atanda et al. 2022; Erbe et al. 2013). Several studies used the genomic prediction approach to harness diversity in germplasm collections (Bari et al. 2021; Crossa et al. 2016; Haile et al. 2020; Jarquin et al. 2016).

Despite the considerable reduction in genotyping costs over the last decade, implementing GS in plant breeding remains challenging for many plant species and breeders due to the need for genotyping the new selection candidates every year and the cost associated to it. As an alternative, Phenomic Selection (PS) has been proposed, replacing molecular information with low-cost phenotypic data, such as Near- InfraRed (NIR) spectra (Rincent et al. 2018; Robert et al. 2022a). NIR spectra have been shown to capture genetic similarities between genotypes, enabling accurate predictions of complex traits.

A key advantage of PS is its low cost, along with its ability to account for non-additive genetic effects, including epistasis and genotype-by-environment interactions (Robert et al. 2022b). Notably, studies have demonstrated that NIR spectra from grains harvested in one environment can be used to predict any complex trait in any environment, as would be done in GS with genotyping (Rincent et al. 2018). In practice, any GS model can be used for PS by simply replacing the marker alleles by the NIR absorbance values at each wavelength after some pre-treatments (Robert et al. 2022a).

The PS has been successfully validated in multiple crops, including bread wheat (Rincent et al. 2018), durum wheat (Meyenberg et al. 2024), rye (Galán et al. 2020), rice (De Verdal et al. 2024), maize (Adak et al. 2023; Rincent et al. 2025), sorghum (Bienvenu et al. 2025), soybean (Zhu et al. 2021), rapeseed (Laurençon et al. 2024; Roscher-Ehrig et al. 2024), grapevine (Brault et al. 2022), and poplar (Rincent et al. 2018), demonstrating predictive abilities often comparable to GS. PS holds particular promise for pea breeding, as NIR spectra are quite often collected in breeding programs to predict protein content. This suggests that in some applications, PS could be implemented at no additional cost. However, to our knowledge, PS has never been tested in pea.

The aims of this study were threefold: (1) to generate a dataset suitable for investigating phenomic selection in peas; (2) to compare the predictive ability of PS, GS and integrated PS-GS models for yield-related traits in elite spring pea varieties; and (3) to evaluate the predictions models for pea breeding improvement.

## Materials and Methods

### Plant materials and field experiments

A panel of 288 spring pea accessions was created by RAGT2n, KWS Momont, UNISIGMA and INRAE. This panel includes 72 public varieties from INRAE CRB PROTEA and 216 breeding lines provided by the three French breeding companies, selected for their agronomic performance and/or their high seed protein content. The panel was phenotyped under twelve environments in field (three years * four locations) in France in 2021, 2022 and 2023 at Estrées-Mons (49°52’41’’N – 03°00’24’’E), Mons-en-Pévèle (50°28’46’’N – 03°06’08’’E), Louville-la-Chenard (48°19’28’’N – 01°47’16’’E) and Froissy (49°34’01’’N – 02°13’16’’E). The 288 lines were cultivated in micro-plots (approximately ∼10m²) in a P- rep design, with 10% of the lines replicated in each environment. Data were collected on the Days To Flowering (DTF, the number of days since 1 January), Thousand Seed Weight (TSW, in grams), seed yield at maturity (Yield, in quintals per hectare), seed protein content at maturity (Protein, as percentage of dry matter), seed protein yield at maturity (Protein Yield, in quintals of protein per hectare), the latter being the product of the seed yield and the seed protein content. However, due to unfavourable environmental conditions, phenotypic data were validated in only 10 to 11 environments. Notably, the Estrèes-Mons site could not be harvested in 2021 due to long rain condition during harvest period that compromise grain development and TSW data was not recorded at the Froissy site in 2022.

### Genotypic data

DNA was extracted from fresh-frozen leaves collected from individual plants representative of the accessions in the panel. Genotyping was conducted using the pea Axiom90K SNP array (Ellis et al. 2023). The raw data were analysed using the Axiom Analysis Suite software, as detailed by Ellis et al. 2023. Only PolyHighResolution SNPs were retained, representing a total of 31,180 SNPs with an average missing data rate of 0.37%. Missing data were then imputed using Beagle (Browning and Browning 2007). A principal coordinate analysis (PCoA) was conducted on Modified Rogers’ distance matrix to visualize the genetic relationships among the 288 genotypes.

### NIRS data

Near-infrared spectroscopy (NIRS) data were acquired at INRAE Dijon for a sample of 200 grams of harvested seeds from each line in each environment. NIRS measurements were carried using a FOSS DS2500, and the analysis was performed using FOSS ISIscan Nova v 10.2.1.13 and FOSS Local configurator v 10.2.1.4 software (FOSS NIRSytems, DK-3400 Hilleroed Danemark). Absorbance was measured from 400 to 2,499.5 nm in 0.5 nm steps. The final spectra covering from the visible to the infrared range are the average of 32 repetitions measured by the spectrometer.

NIRS data were used to predict the seed protein content using a prediction equation developed by INRAE Dijon as described in Lecomte et al. 2023. The predictions were validated by comparing them with reference Kjeldahl values performed on a set of 218 independent samples, representative of the range of protein contents in the panel (R² = 0.942).

Various pre-processing steps were applied to the data to minimize the effects of bias introduced by additive and multiplicative factors in the absorbance measurements. The main objective was to make the measured and true absorbance values more similar for phenomic prediction. This was achieved by calculating the first derivative with a windows size of 131nm data points using the “signal” (Ligges et al. 2015) and “prospectr” (v0.2.0) (Stevens and Ramirez-Lopez 2013) R packages.

### Phenotypic data adjustment and variance component estimation

#### Phenotypic data adjustment

Best Linear Unbiased Estimators (BLUEs, later referred to as adjusted means) were calculated using a linear mixed model that accounts for environmental effects, defined as the combination of year and location. The statistical model was:

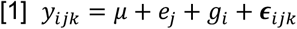

Where 𝑦_𝑖𝑗𝑘_ is the phenotype of genotype i at location j and repetition k. 𝜇 is the overall phenotypic mean, 𝑒_𝑗_ is the fixed effect of location j, 𝑔_𝑖_ is the genotypic fixed effect and 𝜖_𝑖𝑗𝑘_ is the model residual, following a normal distribution 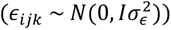. Then, adjusted mean were calculated as *ŷ* = µ + *ĝ* used for the genomic and phenomic prediction models. This adjustment was also used for estimating the BLUEs of first derivatives of each NIRS wavelength from multiple environment NIRS datasets according to the cross-validation scenario and define as *ŝ* (Fig.1). The BLUEs were calculated using R (Version 4.0.5) (R Core Team 2021), with the ’aireml’ package (Version 0.2) (Gilmour et al. 1995).

**Fig. 1.**
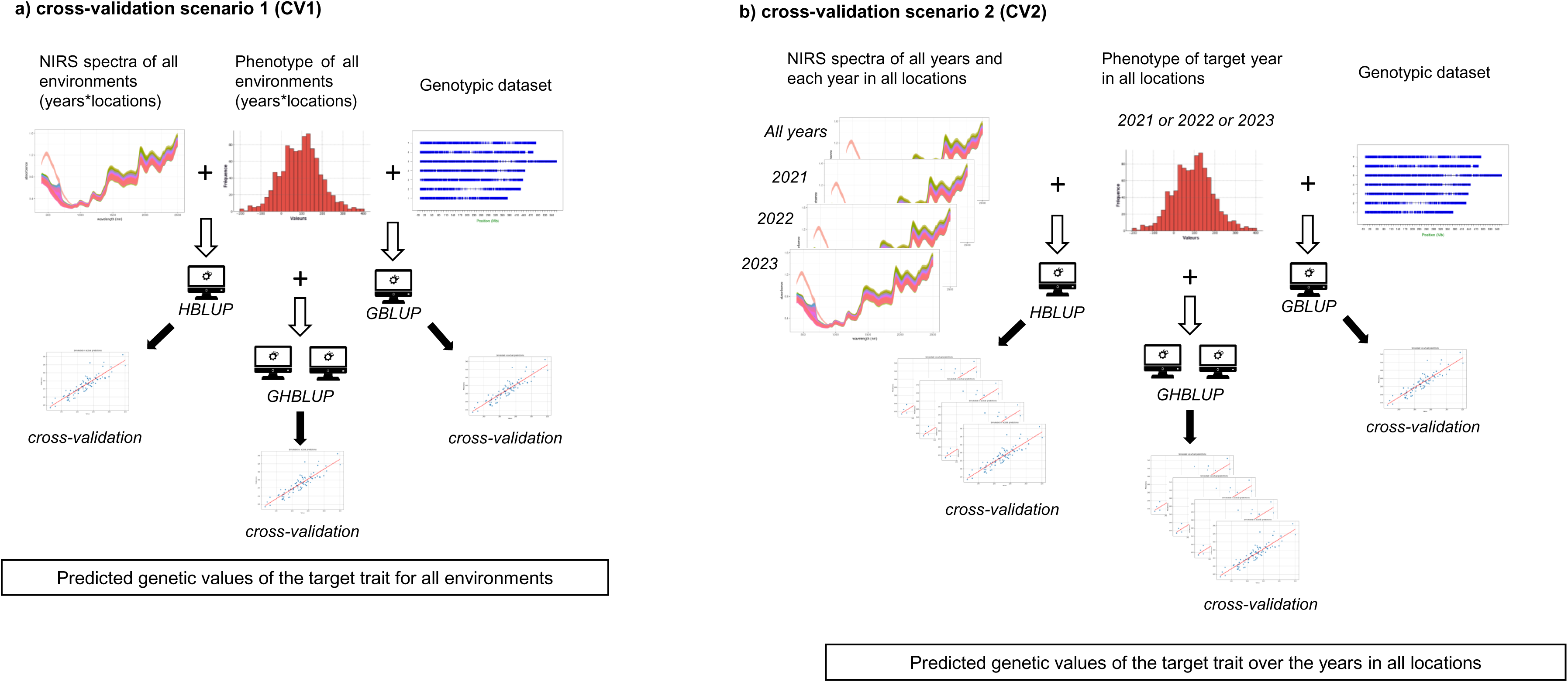
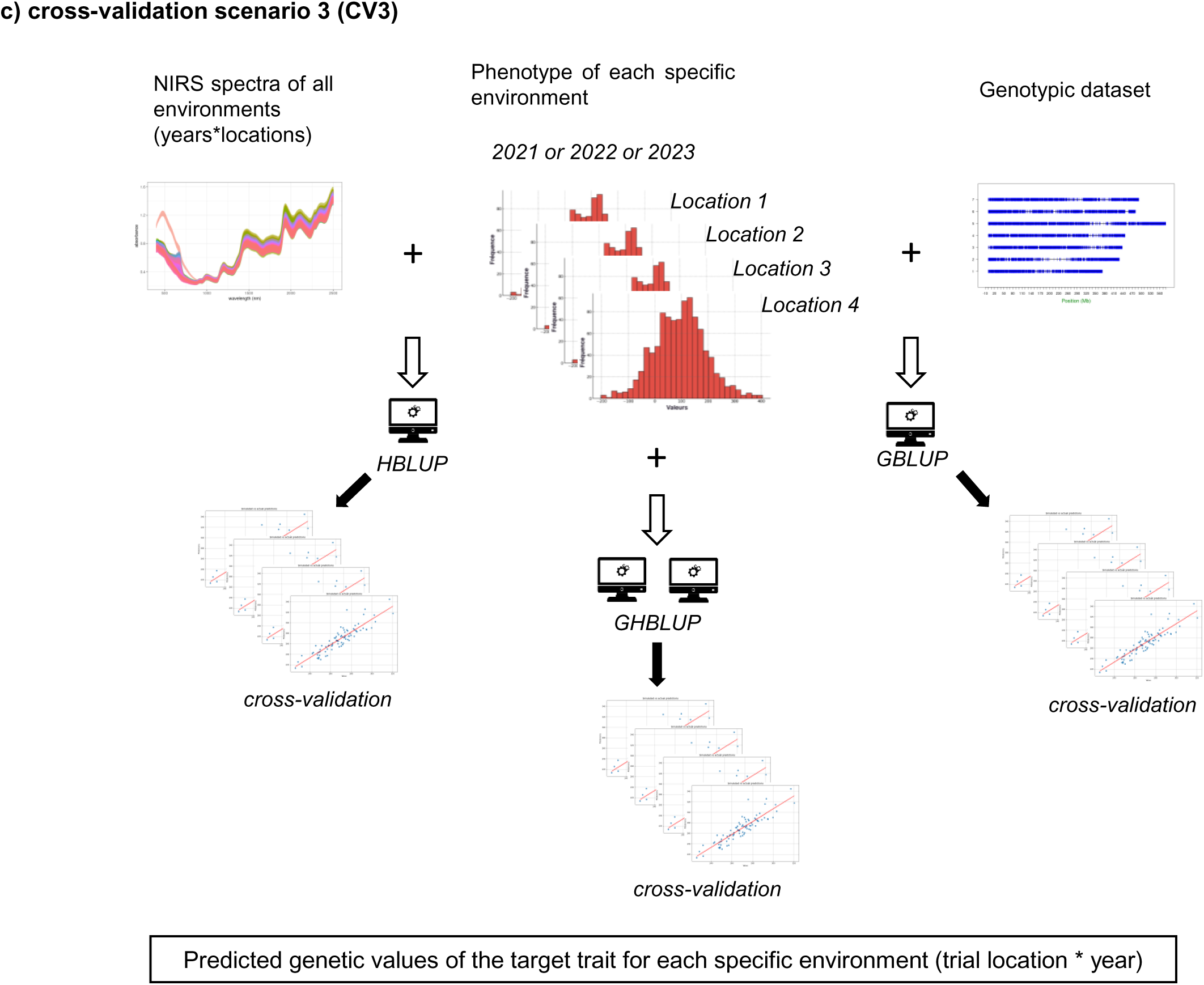
Scenarios of cross-validation (CV1, CV2 and CV3) to performe the predictive accuracy of the HBLUP, GBLUP and GHBLUP models across years and locations. *HBLUP* Hyperspectral Best Linear Unbiased Prediction, *GBLUP* Genomic Best Linear Unbiased Prediction, *GHBLUP* Genomic-Hyperspectral Best Linear Unbiased Prediction

### Variance component estimation

The variances used to calculate trait heritabilities were estimated using the following model:

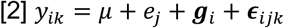

Where 𝑦_𝑖𝑗𝑘_, 𝜇, 𝑒_𝑗_ and 𝜖_𝑖𝑗𝑘_ are defined as in the previous model. 𝒈_𝑖_ is the genetic value, modelled as a random effect following a normal distribution 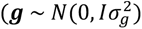. Broad sense heritability was computed as:

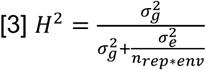

where 𝑛_𝑒𝑛𝑣_ is the number of environments in the multi-environmental trial (combination of 3 years and 4 locations = 12), and 𝑛_𝑟𝑒𝑝∗𝑒𝑛𝑣_ = 13.2 is the average number of repetitions computed as the multiplication of the number of repetitions per environment (1.1) by the number of environments.

### Genomic and Phenomic prediction models

Our objective was to compare the accuracy of a classical genomic prediction model (GBLUP), a phenomic prediction model (HBLUP) and a model combining (GHBLUP) (Convex combination) of the genomic and phenomic (NIRS) information.

The three mixed models were implemented using R (Version 4.0.5) (R Core Team 2021), with the ’cpgen’ package (Version 0.2) (Heuer 2016).

### GBLUP: Genomic Best Linear Unbiased Prediction

The GBLUP model predicts the additive genetic value of genotypes on the basis of their genomic additive relationship. We used a realized additive genomic relationship matrix described in VanRaden (2008):

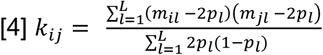

Where, 𝑚_𝑖_ is the allele dose of genotype i for SNP 𝑙 (0/1/2). 2𝑝_𝑙_ and 𝑝_𝑙_ are respectively the mean allele dose and the allele frequency for SNP 𝑙. L is the total number of SNPs.

The GBLUP model is defined as follows:

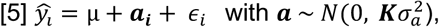

Where 𝒚 is the vector of adjusted means (BLUEs), µ is the overall phenotypic mean. 𝑎_𝑖_ is the random genetic effect following the distribution 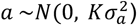. 𝝐 is the error term 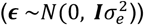

### HBLUP: Hyperspectral Best Linear Unbiased Prediction

The HBLUP model predicts a genetic value on the basis of their NIRS-based relationship, estimated as in Rincent et al. (2018):

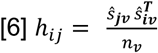

*ŝ*_𝒗_ are the BLUEs of the first derivative of the absorbance at wavelength 𝑣

The HBLUP model is defined as follows:

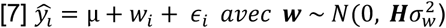

here 𝒚 is the vector of adjusted means (BLUEs), µ is the overall phenotypic mean. 𝑤_𝑖_ is the genetic value following the distribution 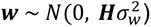. 𝝐 is the vector of residuals 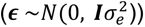.

### GHBLUP: Genomic-Hyperspectral Best Linear Unbiased Prediction

The GHBLUP model predicts a genetic effect based on the convex combination of the genomic relationship matrix **K** and the NIRS-based relationship matrix **H** (Schaid 2010; Smits and Jordaan 2002):

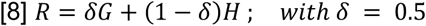

The GHBLUP model is defined as follows:

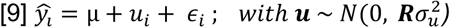

In this model, 𝑢_𝑖_ is the genetic value following the distribution 𝑦_𝑖_ = µ + 𝑢_𝑖_ + 𝜖_𝑖_ ; 𝑤𝑖𝑡ℎ 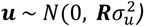. 𝝐 is the vector of residuals 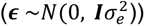.

### Evaluation of prediction accuracy using cross-validation

To evaluate and compare the predictive performance of the GBLUP, HBLUP, and GHBLUP models, we implemented a structured cross-validation framework. Prediction accuracy was consistently quantified as the Pearson correlation coefficient (r) calculated between the predicted genetic values derived from each model and the observed phenotypic BLUEs within the respective testing sets. To ensure the stability and reliability of our accuracy estimates, each cross-validation procedure outlined below was repeated 10 times using different random partitions of the data, and the average correlation across these repetitions was reported as the final accuracy metric.

For all scenario, a 5-fold cross-validation scheme was applied to this aggregated dataset. Within each of the 10 repetitions, genotypes were randomly assigned to one of five equally sized folds. Models were then iteratively trained using data from four folds (representing 80% of the genotypes, the training set) and subsequently validated on the remaining fold (the testing set, comprising 20% of the genotypes). Overall each cross-validation scheme contains 50 models.

The models evaluated under this global scenario included GBLUP, HBLUP, and the combined GHBLUP models. We used three distinct cross-validation scenarios across years and locations. These cross-validation scenarios are visually depicted in Fig.1.

### Cross-Validation 1 (CV1)

The first scenario, designated as global prediction performance assessment, aimed to evaluate the predictive ability of the models when trained on data from all the environments tested in the present study. This approach simulated the challenge of predicting genotype performance across the entire target population of environments, thereby assessing the generalizability of the models (Fig.1a).

### Cross-Validation 2 (CV2)

The second scenario assessed the predictive abilities of models trained on specific years to understand the robustness of predictions over the years and the impact of year-specific environmental conditions (Fig.1b). For this purpose, BLUEs computed for each year over the field-trial sites. This stratification resulted in three independent subsets, corresponding to the data of 2021, 2022, and 2023. Four NIRS datasets were used to train HBLUP and GHBLUP to predict the specific years: “All” NIRS combining all data, and NIRS recorded from field testing in 2021, 2022 and 2023. This scenario allowed the detailed comparison of prediction models when both training and testing datasets originated from the same year. It specifically allowed for an examination of the relative merits of using global NIRS information versus yearly NIRS data for prediction within a given year (Fig.1b).

### Cross-Validation 3 (CV3)

The third scenario evaluated the prediction performance within each specific environment, defined by the trial location and year (Fig.1c). This approach allowed assessing the performance of environment specific models, that are expected to better reflect the environmental conditions and potential genotype-by-environment interactions specific to an environment. To implement this, BLUEs were computed for each single environment.

## Results

### Phenotypic variability

DTF, TSW, Yield, Protein, and Protein Yield were measured on a panel of 288 lines between 2021 and 2023 in field trials network at four locations (Table 1). Phenotypic data could be validated in 10 to 11 environments (location x year) per trait out of 12 environments. Mean of DTF was 153.59 days with a low variation between lines (sd=10.45, CV=0.07). DTF exhibited high heritability (h²=0.93), indicating strong genetic control. The average of TSW is 231.88 g, with a greater variation between lines (sd= 30.39, CV= 0.13) and a high heritability (h²=0.94). Heritability was also high for protein content (h=0.91) with an average of 24.05% and very low variation between lines (sd=2.03, CV=0.08). In contrast, yield (mean= 43.31 qx/ha) and protein yield (mean= 18.15 qx/ha) displayed greate phenotypic variability, with coefficients of variation of 0.30 and 0.32, respectively. These traits had comparatively lower heritability values, estimated at 0.84 for yield and 0.82 for protein yield.

**Table 1.** Descriptive statistics of phenotypic data collected from a field trials network comprising fours locations each year between 2021 and 2023

| Trait<sup>a</sup> | NbData<sup>b</sup> | NbInd<sup>c</sup> | NbEnv<sup>d</sup> | mean | sd<sup>e</sup> | CV<sup>f</sup> | Heritability |
| --- | --- | --- | --- | --- | --- | --- | --- |
| DTF (date) | 3501 | 288 | 11 | 153.59 | 10.45 | 0.07 | 0.93 |
| TSW (g) | 3170 | 288 | 10 | 231.88 | 30.39 | 0.13 | 0.94 |
| Yield (quintals/hectare) | 3494 | 288 | 11 | 43.31 | 12.96 | 0.30 | 0.84 |
| Protein (%) | 3491 | 288 | 11 | 24.05 | 2.03 | 0.08 | 0.91 |
| Protein Yield (quintals of protein/hectare) | 3490 | 288 | 11 | 18.15 | 5.72 | 0.32 | 0.82 |

<sup>a</sup> *DTF* Days To Flowering (date), *TSW* Thousand Seed Weight (gram), *Yield* seed yield at maturity (quintals per hectare), *Protein* seed protein content at maturity (percentage of dry matter), *Protein Yield* seed protein yield at maturity (quintals of protein per hectare)
<sup>b</sup> Number of data, <sup>c</sup> Number of individuals, <sup>d</sup> Number of environments, <sup>e</sup> Standard deviation, <sup>f</sup> Coefficient of variation

Phenotype distributions and pearson correlation coefficients between phenotypic traits over all environments (Fig.2) showed that DTF was significantly and negatively correlated with TSW (r= - 0.22) and Protein content (r= -0.12), but showed no significant correlation with the Yield (r= -0.073). Conversely, TSW exhibited significant and positive correlation with Yield (r=0.29) and Protein_Yield (r=0.26), indicating its contribution to overall productivity. As expected, Yield was significantly and positively correlated with Protein Yield (r=0.93), but negatively correlated with Protein content (r= - 0.36) highlighting a trade-off between productivity and seed quality in this panel.

**Fig. 2.**
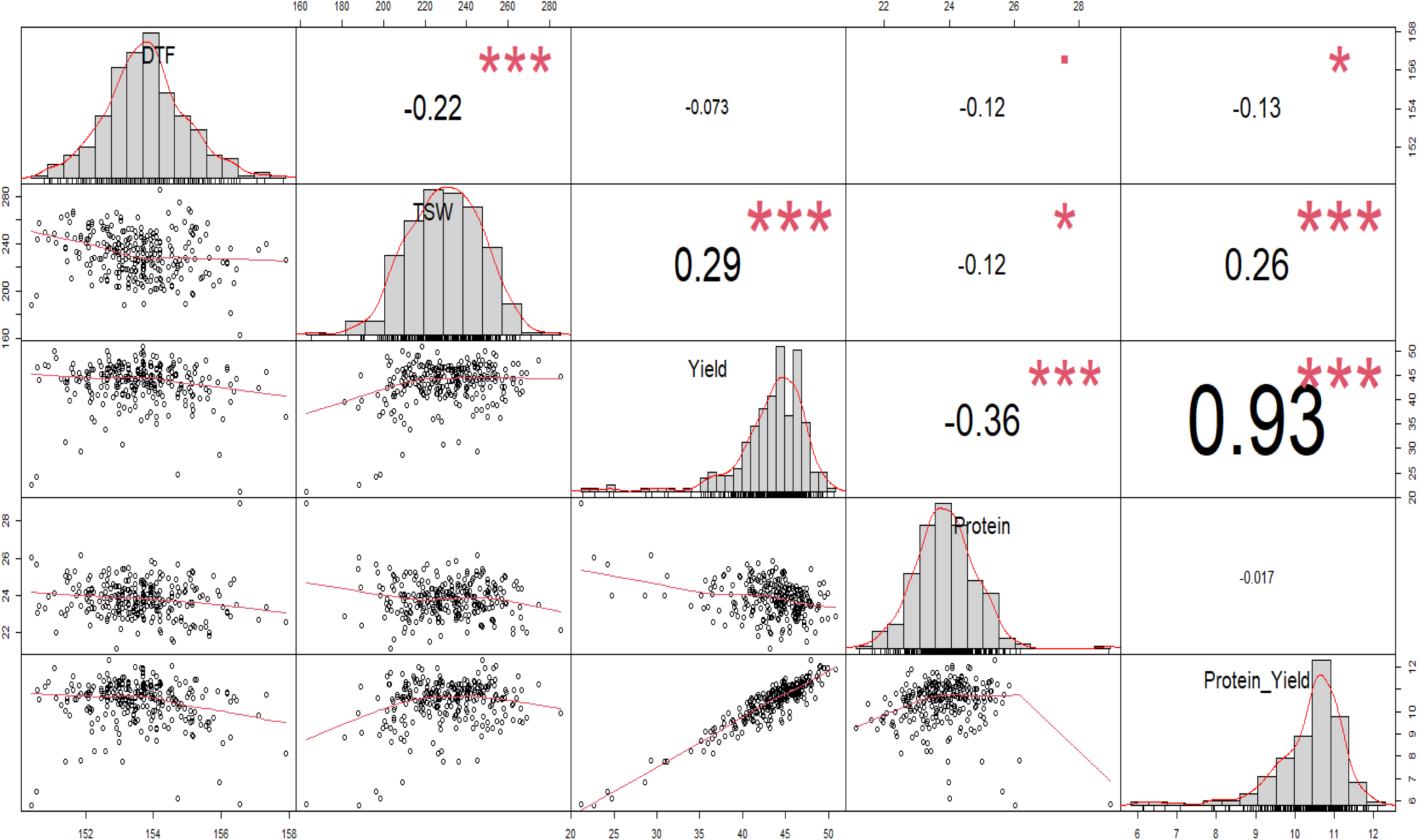
Pearson correlation coefficients betwenn phenotypic traits recorded from 2021 to 2023 in a field trials network of fours locations per growing season . *DTF* days to flowering (date), *TSW* thousand seed weight (g), *Yield* seed yield at maturity (quintals/ha), *Protein* seed protein content at maturity (% of dry matter), *Protein_Yield* seed protein yield at maturity (quintals of protein/ha). *, ** and *** significant correlation at the P<0.05, P<0.01 and P<0.001 probability level, respectively

The phenotypes’ distributions by environments observed from 2021 to 2023 are illustrated in the supplementary Fig.S1. The datasets are available from the corresponding author on reasonable request.

### Genotypic diversity

The Axiom90K SNP array was used to genotype the 288 spring pea accessions under investigation and a comprehensive dataset of 31,180 high-quality markers was obtained. The initial analysis revealed that 0.37% of the data was missing and imputation was undertaken using Beagle software. The imputed matrix was used for the diversity analysis. The distribution of the markers was fairly homogeneous, with an average of 4,942 SNPs per chromosome, a maximum for chromosome 5 with 7,012 SNPs and a minimum for chromosome 2 with 3,127 SNPs. Average distance between markers is 0.03±0.04 Mb (supplementary Fig.S2). An important part of the SNPs showed low Minor Allele Frequency (MAF), but there are more than 20K SNPs with MAF >10% (supplementary Fig.S3). The first two axes of the PCoA showed some clustering between partners’ lines (Fig.3). Partner 1 lines being more distant than lines from partner 2 and 3. The public varieties assured an overlap between the three breeder lines’ sets. This overlap is likely due to breeders using genetically similar cultivars as parents in crosses, while divergence or different groups within partners could be explained by the fact that each breeder having their own source of parental lines. The overlap appears large enough that a specific cross-validation schemes is not required to account for genetic diversity.

**Fig. 3.**
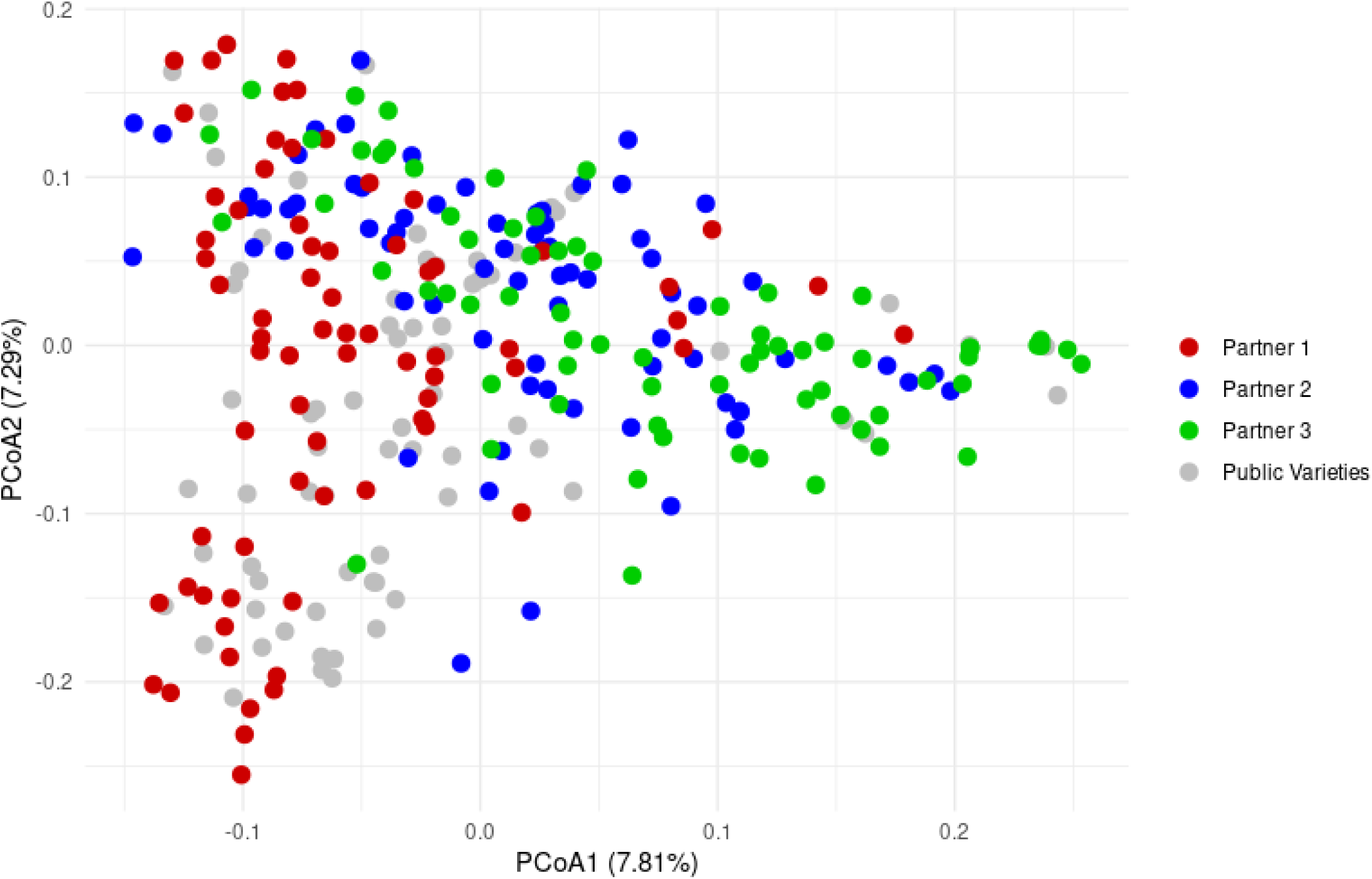
Population structure analyses of the 288-accession spring pea panel using a multidimensional principal coordinate analysis (*PCoA*) with 31,180 high-quality SNP markers. Points on the graph, representing accessions, are color-coded based on donors. Red, Blue and green refer to breeding lines from partner 1, 2 or 3, respectively. Current and old varieties are in grey

### Variance of NIR spectra

The NIRS absorbance spectra figure, as shown in supplementary Fig.S4, displays the absorbance values (y-axis) across a range of wavelengths from 400 nm to 2,499.5 nm (x-axis). Each colored line represents the raw spectrum for a different pea line. The spectra exhibit characteristic peaks and troughs, indicating wavelengths where light is more or less absorbed by the grain samples. For instance, there are notable peaks in absorbance around 1,200 nm, 1,450-1,500 nm, 1,700-1,800 nm, 1,900-2,000 nm, and broader absorbance features in the 2,200-2,499.5 nm region. The spread of the lines across the absorbance values at any given wavelength indicates variability among the samples. A particularly high variability among samples appears in the visible region (around 400-700 nm) and in some specific NIR regions, such as the peak around 1,200 nm and the region above 1,900 nm. This variability is what phenomic selection aims to exploit, as these spectral differences can be correlated with genetic and phenotypic traits.

Total variance partitioning of NIRS spectra obtained from seeds harvested from different accessions and locations (see Fig.4) shows that the environmental component (shown in green) accounts for the largest proportion of variance across most of the spectrum (400–2,499.5 nm), particularly between 700 and 2,499.5 nm. However, the genotypic component (red) shows significant contributions in specific regions. Prominent peaks for genotypic variance are evident in the visible range (approximately 400– 700 nm), particularly around 600–700 nm (green and yellow light). Distinct yet smaller peaks are also present in the NIR region, for example at around 1,100–1,200 nm, 1,400 nm, and 2,300 nm. The residual variance (blue) is generally the smallest component, but it becomes more substantial at the extremes of the measured range and in some narrow bands. Regions of the visible light spectrum with higher genotypic variance are likely due to the presence of differently coloured grains (green and yellow). Nevertheless, the remaining genetic variances are of interest for phenomic selection as they indicate wavelengths at which spectral data is more strongly influenced by genetic differences between pea lines.

**Fig. 4.**
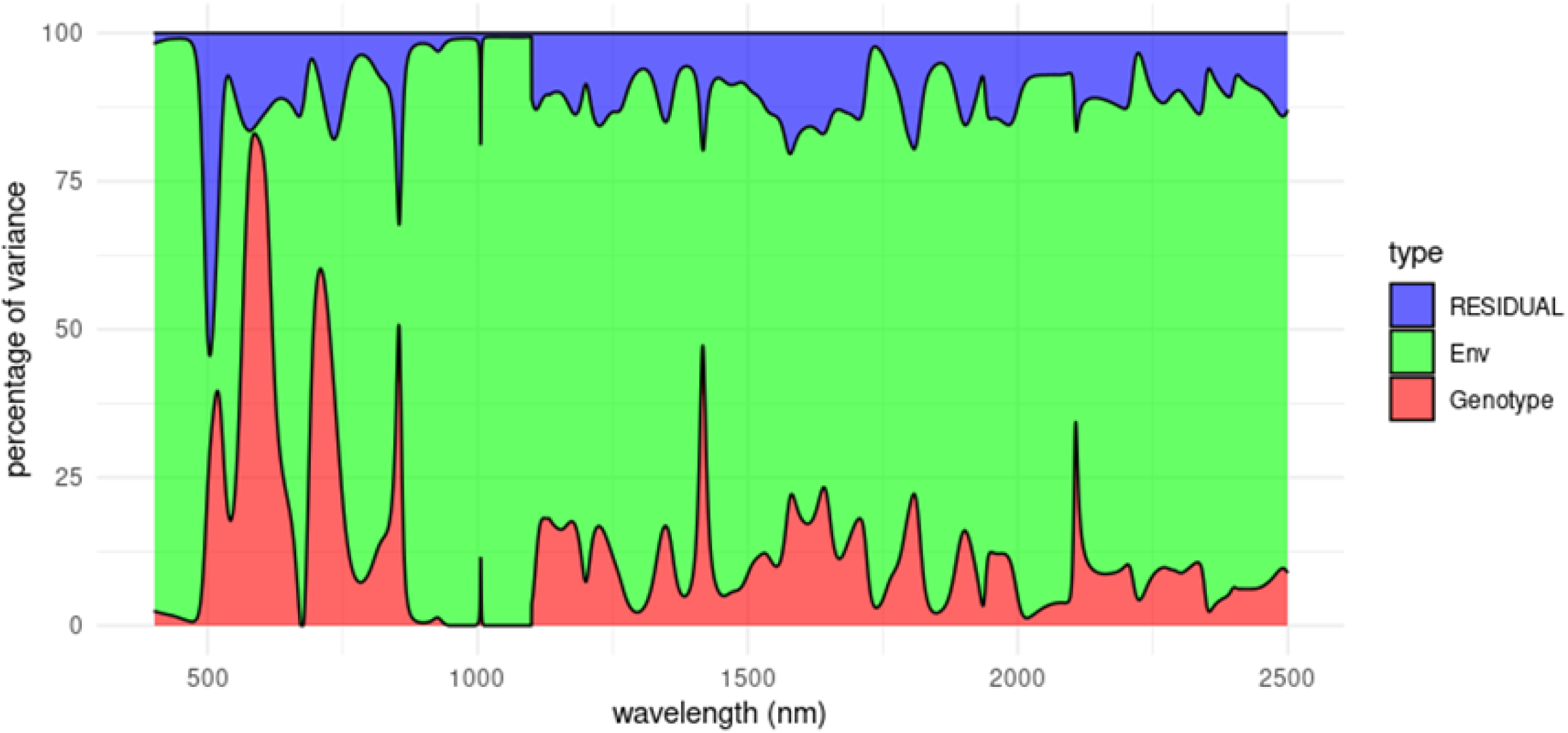
Total variance partitioning of NIRS spectra from the seeds of the 288-accession panel, collected from all validated environments. The x-axis represents the wavelengths considered (400 nm to 2,499.5 nm), while the y-axis shows the percentage of total NIRS absorbance variance that can be attributed to genotype (red), environment (green) and residual (blue) factors across the measured wavelengths

### Model prediction accuracy

The predictive accuracy of genomic, phenomic, and combined genomic-phenomic models was assessed for five key agronomic traits measured: Days To Flowering (DTF), Thousand Seed Weight (TSW), seed yield, seed protein content and seed protein yield. Prediction accuracy is expressed as the mean Pearson correlation between the observed values and predicted scores obtained through cross-validation in one of three scenarios: CV1, CV2 and CV3 (see Fig.1a,b,c).

In the first scenario (CV1, Fig.1a), the genomic prediction model exhibited the highest predictive accuracy for DTF (0.56), followed by the combined genomic-phenomic model (0.53) and the phenomic model (0.45) (Fig. 5). The inclusion of phenomic data did not improve the prediction of this trait. In contrast, protein content prediction was significantly improved using phenomic data (0.91), and was further enhanced by the combined model (0.94), both outperforming the genomic model alone (0.65). For protein yield, the combined model considering genomic and phenomic data achieved the highest accuracy (0.81), compared to the phenomic (0.74) and genomic (0.62) models, highlighting the benefit of integrating both data sources. Regarding TSW, prediction accuracies were relatively similar across all models: 0.74 for the phenomic, 0.72 for the combined model and 0.70 for the genomic model. Finally, yield was most accurately predicted by the combined model (0.76), outperforming the genomic (0.67) and phenomic (0.62) models highlighting the advantage of combining both data sources for complex trait prediction.

**Fig. 5.**
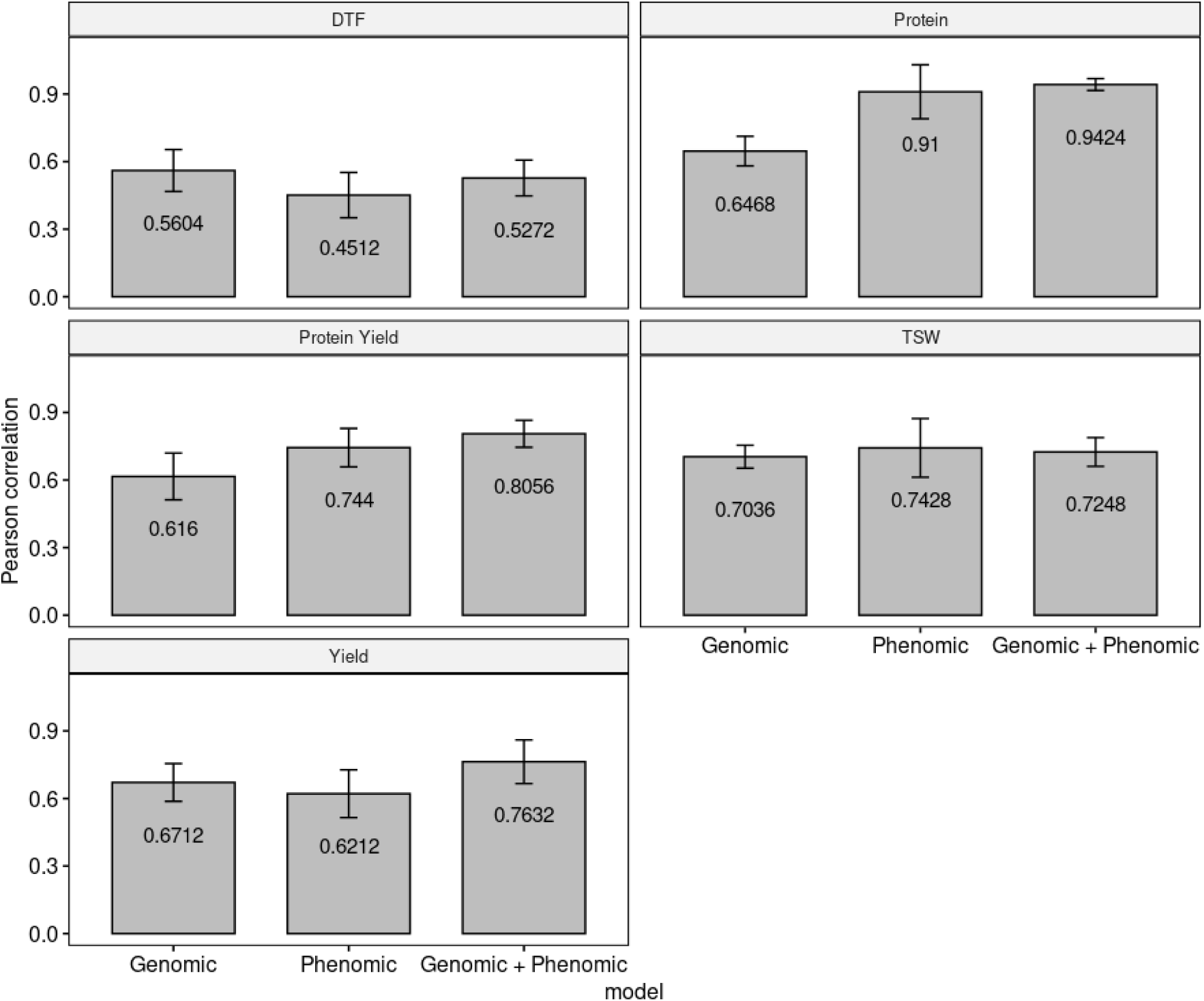
Predictive accuracy of Genomic, Phenomic and combined Genomic-Phenomic models in CV1 scenario. Prediction accuracies are expressed as Pearson coefficients correlations between observed and predicted values for Day To Flowering (DTF), Protein, Protein Yield, Thousand Seed Weight (TSW) and seed Yield. The bars indicate the standard deviations of the ten replicates of the five-fold cross-validations

In the second cross-validation scenario (CV2, Fig.1b), which assesses the predictive performance of the models across individual years, prediction accuracy varied notably depending on the year used to train the model (Table 2). Overall, using NIRS data from year *n* resulted in the highest predictive accuracy for phenotypes measured in that same year. For instance, phenotypes of 2021 were best predicted using the 2021 spectral data to train a model (mean accuracy = 0.63), outperforming models that used spectra from 2022 (0.52), 2023 (0.48), or from all three years combined (0.58). Genomic prediction alone achieved a mean accuracy of 0.57 for the same year. This pattern was similarly observed in 2022, where the highest prediction accuracy (0.67) was achieved using 2022 NIRS data. In contrast, for 2023, the best performance was obtained using the combined spectra from all three years (mean = 0.62). Across all years, the integration of genomic and phenomic data consistently resulted in higher predictive accuracies compared to using either data source alone. This improvement was particularly notable for complex traits such as yield, for which accuracy increased by approximately 10%. Importantly, the optimal configuration for phenotype prediction within a given year consistently involved the use of genomic data in combination with NIRS data from the same year.

**Table 2.** Predictive accuracy of Genomic, Phenomic and combined Genomic-Phenomic models in CV2 scenario. Prediction accuracies are expressed as Pearson coefficients correlations between observed and predicted values for each year in all locations for Day To Flowering (DTF), seed Protein, seed Protein Yield, Thousand Seed Weight (TSW) and seed Yield. Best accuracy are in grey

|  | 2021 (3 environments) |  |  |  |  |  |  |  |  |
| --- | --- | --- | --- | --- | --- | --- | --- | --- | --- |
|  | Genomic | Phenomic |  |  |  | Genomic + Phenomic |  |  |  |
|  |  | All | 2021 | 2022 | 2023 | All | 2021 | 2022 | 2023 |
| DTF | 0.50 | 0.33 | 0.34 | 0.39 | 0.50 | 0.47 | 0.51 | 0.50 | 0.55 |
| Protein | 0.50 | 0.82 | 0.91 | 0.67 | 0.58 | 0.79 | 0.88 | 0.70 | 0.64 |
| Protein Yield | 0.58 | 0.61 | 0.64 | 0.54 | 0.48 | 0.69 | 0.69 | 0.66 | 0.66 |
| TSW | 0.70 | 0.63 | 0.70 | 0.53 | 0.46 | 0.71 | 0.70 | 0.72 | 0.68 |
| Yield | 0.58 | 0.51 | 0.56 | 0.45 | 0.39 | 0.62 | 0.66 | 0.62 | 0.61 |
| mean | 0.57 | 0.58 | 0.63 | 0.52 | 0.48 | 0.66 | 0.69 | 0.64 | 0.63 |

|  | 2022 (4 environments) |  |  |  |  |  |  |  |  |
| --- | --- | --- | --- | --- | --- | --- | --- | --- | --- |
|  | Genomic | Phenomic |  |  |  | Genomic + Phenomic |  |  |  |
|  |  | All | 2021 | 2022 | 2023 | All | 2021 | 2022 | 2023 |
| DTF | 0.52 | 0.48 | 0.43 | 0.50 | 0.51 | 0.51 | 0.53 | 0.55 | 0.59 |
| Protein | 0.59 | 0.86 | 0.71 | 0.91 | 0.68 | 0.86 | 0.74 | 0.91 | 0.76 |
| Protein Yield | 0.52 | 0.69 | 0.62 | 0.67 | 0.58 | 0.74 | 0.68 | 0.73 | 0.68 |
| TSW | 0.63 | 0.69 | 0.66 | 0.72 | 0.52 | 0.71 | 0.68 | 0.72 | 0.66 |
| Yield | 0.55 | 0.60 | 0.53 | 0.56 | 0.48 | 0.65 | 0.66 | 0.70 | 0.63 |
| mean | 0.56 | 0.66 | 0.59 | 0.67 | 0.55 | 0.70 | 0.66 | 0.72 | 0.66 |

|  | 2023 (4 environments) |  |  |  |  |  |  |  |  |
| --- | --- | --- | --- | --- | --- | --- | --- | --- | --- |
|  | Genomic | Phenomic |  |  |  | Genomic + Phenomic |  |  |  |
|  |  | All | 2021 | 2022 | 2023 | All | 2021 | 2022 | 2023 |
| DTF | 0.58 | 0.40 | 0.35 | 0.42 | 0.41 | 0.54 | 0.58 | 0.57 | 0.58 |
| Protein | 0.68 | 0.84 | 0.68 | 0.81 | 0.86 | 0.89 | 0.81 | 0.87 | 0.91 |
| Protein Yield | 0.58 | 0.64 | 0.50 | 0.64 | 0.65 | 0.71 | 0.64 | 0.70 | 0.72 |
| TSW | 0.69 | 0.69 | 0.61 | 0.63 | 0.62 | 0.69 | 0.67 | 0.71 | 0.71 |
| Yield | 0.57 | 0.54 | 0.42 | 0.55 | 0.53 | 0.62 | 0.58 | 0.66 | 0.64 |
| mean | 0.62 | 0.62 | 0.51 | 0.61 | 0.61 | 0.69 | 0.66 | 0.70 | 0.71 |

When predicting a particular trial (CV3, Fig. 1c), genomic and phenomic predictive abilities were variable from one trial to another (Table 3). For yield, the highest predictive ability was equal to 0.55 for genomic prediction at Froissy in 2022, and the lowest was equal to 0.27 at Louville-la-Chenard in 2023. For the same trait, the phenomic predictive abilities varied from 0.23 (Louville-la-Chenard) to 0.59 (Froissy 2022). On average across all trials, genomic prediction was more accurate (0.46) than phenomic prediction (0.41) for yield, but the best model was the one combining genomic and phenomic (0.52). On average, phenomic predictions were more accurate or similar to genomic predictions for protein, protein yield, and TSW, while genomic predictions were more accurate for DTF. Most importantly, the model combining genomic and phenomic was better than phenomic or genomic alone (except for DTF for which it was similar to genomic prediction). On average across all trials and all traits, genomic and phenomic predictions reached predictive abilities of 0.49 and 0.50, respectively, while the model combining both sources of information reached 0.57.

**Table 3.**
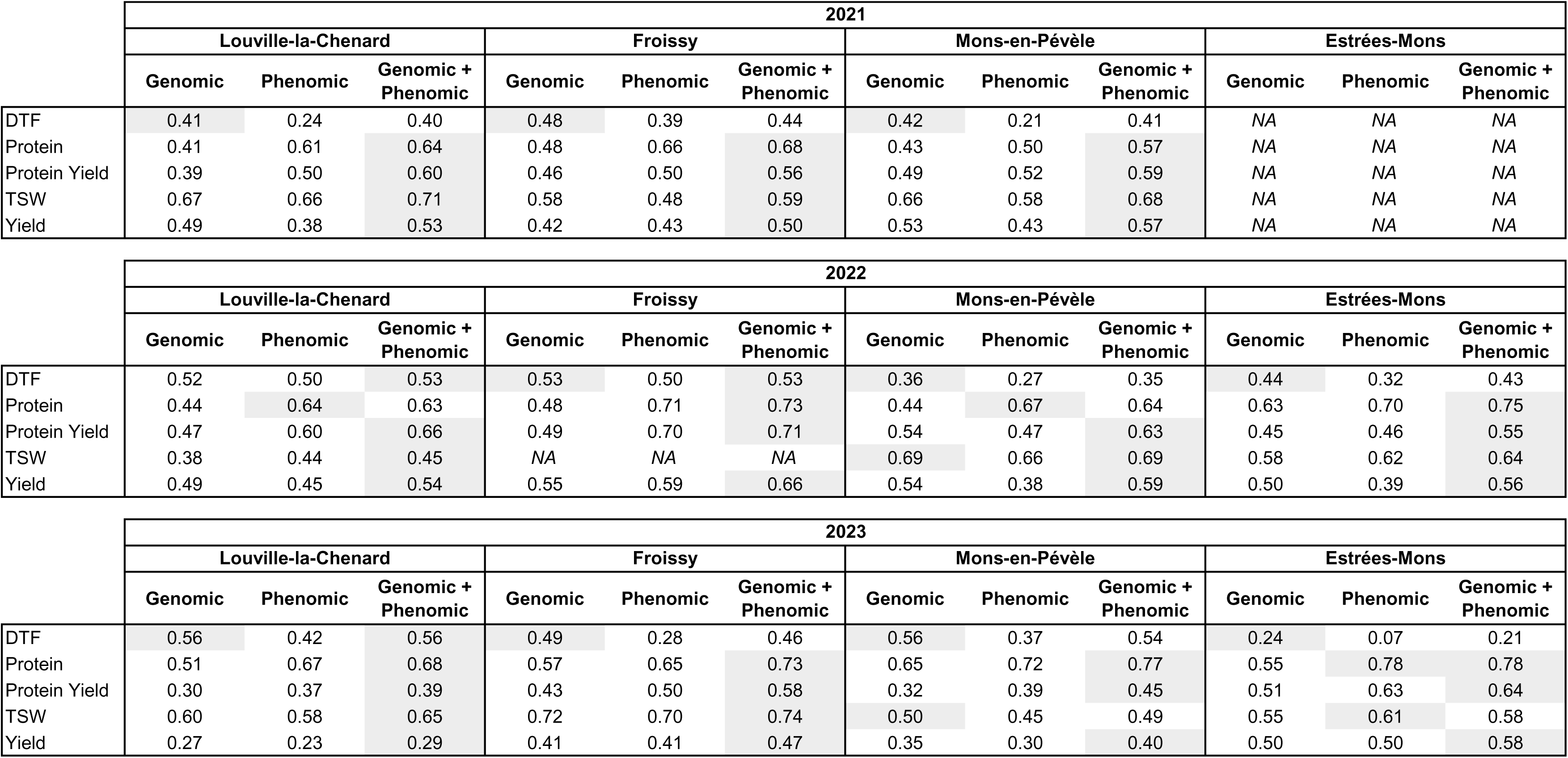
Predictive accuracy of Genomic, Phenomic and combined Genomic-Phenomic models in CV3 scenario. Prediction accuracies are expressed as Pearson coefficients correlations between observed and predicted values for each year in each location for Day To Flowering (DTF), seed Protein, seed Protein Yield, Thousand Seed Weight (TSW) and seed Yield. *NA* missing values. Best accuracy are in grey

## Discussion

### Structuration of data set

The primary objective of this study was to establish a dataset enabling the investigation of genomic and phenomic selection within pea breeding programs. The diversity panel comprised 288 spring pea accessions, including cultivars (registered lines) and advanced breeding lines contributed by three independent French breeding companies. Breeders were encouraged to propose genetically diverse materials with contrasting phenotype, particularly for seed yield and seed protein content, two traits of major breeding interest. This strategy led to the inclusion of germplasm reflecting a broad range of genetic backgrounds and breeding objectives, as each partner applied independent selection schemes and trait priorities. Although the pea benefit from relatively open germplasm exchange among breeding programs (Magrini et al. 2016), the analysis revealed some clustering of the genetic structure within the panel. This clustering may influence the predictive accuracy (Crossa et al. 2017). However, the genetic overlap observed among the partner materials herein suggests that cross-validation predictions across breeding programs may be feasible. Nevertheless, this genetic overlap may also lead to an overestimation of genetic variances and prediction accuracies in cross-validation frameworks, especially under scenarios where the training and validation sets are not fully independent (Isidro et al. 2015). The relatively balanced representation of diverse genetic backgrounds could artificially inflate predictive metrics compared to what might be observed in more homogeneous or stratified breeding materials. Consequently, the predictive performance observed in this study may exceed what is typically achievable in more genetically uniform breeding populations.

Importantly, the traits analyzed in this study exhibited high heritability estimates, indicating strong genetic determinism and supporting the feasibility of both genomic and phenomic prediction approaches. At the same time, high coefficients of variation for key traits such as yield and protein yield underscore the influence of environmental variability and the challenge of achieving stable trait performance across diverse growing conditions. These findings further highlight the necessity of incorporating genotype-by-environment (G×E) interactions into selection models to improve robustness (Jarquín et al. 2014). The phenotypic correlation structure observed in our dataset also illustrates the inherent challenge of simultaneously improving yield and seed quality in pea breeding. In particular, the moderate negative correlation between yield and seed protein content reflects the classical trade-off observed in breeding. However, traits such as thousand seed weight (TSW) and protein yield emerge as promising targets for balanced selection strategies (Michel et al. 2019). The positive correlations between TSW and yield, as well as TSW and protein yield, suggest that TSW could serve as an effective indirect selection criterion for enhancing overall productivity without compromising quality. These results support the integration of multi-trait selection models to optimize genetic gains for both yield and quality traits, as recently demonstrated by Saludares et al. (2024) in field pea trials.

### Predictive ability

As shown in other studies, our results confirm that NIR spectra collected within a given year generally provide more accurate predictions for that same year compared to using spectra from different years. This can be explained by the ability of the spectra to capture genotype-by-environment (GxE) interactions (Robert et al. 2022b), but could also be due to environmental effects leading to overoptimistic accuracies (Wang et al. 2025). However, in our dataset, this effect seem marginal, with only minor differences in predictive accuracy between years when predicting a given year.

A key conclusion of this study, is that the spectra acquired in one year can be used to accurately predict in other years, confirming the potentiel use of NIRS to predict traits independent from the tissue on which the spectra are acquired (Rincent et al. 2018). This opens the way to promising applications of phenomic selection, with for instance the possibility to acquire NIR spectra in nurseries to predict productivity or other complex traits that are unavailable at this step of the breeding program.

In this study, we explored three predictive scenarios to assess model efficiency under contrasting conditions. An additional promising scenario, not explored here, is sparse testing. In this approach, genotypes are observed in only a subset of environments, raising the challenge of integrating spectral data from different environments with missing genotype-environment combinations (Robert et al. 2022b). However, the implementation of sparse testing strategies enables efficient phenotyping with negligible compromise in predictive ability (Garcia-Abadillo et al. 2024), representing a major advantage for large-scale breeding programs. Furthermore, although we used simple kernel-based models, alternative approaches such as machine learning and deep learning could capture complex nonlinear effects and interactions between SNPs, wavelengths, and between SNPs and wavelengths, for instance as proposed by Wang et al. (2023) for multi-omics integration. The advanced modeling methods could extend the predictive ability in the unknown environments (Paleari et al. 2025).

### Integrate predictive models in breeding programs

The evaluation of predictive models revealed distinct patterns regarding their effectiveness. The genomic selection approach provided moderate to good predictive accuracy, particularly for yield-related traits. However, its capacity to fully capture the phenotypic variability expressed across multiple environments appeared limited, reflecting the challenges of accounting for genotype-by-environment interactions through genomic information alone (Teressa et al.2021). In contrast, phenomic model based on spectral data demonstrated strong predictive accuracy for specific traits, such as protein yield. These results highlight the ability of phenomic selection to improve strategic trait for pea breeding selection, consistent with results observed in other crop species (Adunola et al. 2024; Dallinger et al. 2023; DeSalvio et al. 2024). Nonetheless, they were less effective in predicting phenological traits like flowering time, likely due to the variability of these traits under variable environmental conditions.

The integration of phenomic and genomic data in an integrated predictive model consistently led to improved predictive performance across all traits except for the flowering. This integrative approach combines the heritable information captured by molecular markers, and environment-responsive signals provided by phenomic measurements. This synergy was particularly evident in the enhanced prediction of complex traits such as protein yield and overall grain yield and offers notable improvements in selection and pea breeding programs. These results support current trends in data-assisted breeding and confirms that multi-source predictive models are essential for improving the robustness, accuracy, and transferability of predictions (Graciano et al. 2025; Sandhu et al. 2021). By combining molecular and spectral insights, such integrative strategies provide a powerful framework to accelerate genetic gain in pea breeding programs, especially under increasingly variable agro-environmental conditions.

## Author Contribution Statement

All authors contributed to the study. AK wrote the manuscript, performed statistical analysis and managed the plant material, VF performed the statistical analysis, AW performed the statistical analysis and contributed to the manuscript writing, GA managed and analysed the genotypic data, MNH, MT and JR provided the NIRS data, JFH and GR developed the plant material and produce the phenotypic data, CK and ACL developed the plant material, NT and RR contributed to the manuscript writing, PD and JB initiated the study, GT supervised the study and contributed to the manuscript writing. All authors read and approved the final manuscript.

## Acknowledgments

The authors thank the work in experimental units of INRAE Estrèes-Mons, RAGT2n, Unisigma and KWS Momont for contributing to field experiments. The authors thank the GENTYANE platform for genotyping. The authors thank the pulse genetic resource centre managed by INRAE (CRB PROTEA), particularly Marianne Chabert-Martinello and Mathieu Chanis for the supply of public pea accessions. We acknowledge Audrey Courtial, Karen Boucherot and Catherine Desmetz for the implication in the genotyping. We acknowledge Hervé Houtin for managing seed collection and distribution and, Florian Barthes and Eric Hanocq for the contribution in the field experiments and the phenotyping.

## Availability of data and material

The datasets generated during and/or analysed during the current study are not publicly available due to breeding programs privacy but are available from the corresponding author on reasonable request. Code used to lead the analysis of this study is available from the corresponding author on request.

## Ethical approval

The experiments comply with the current laws of the country in which they were performed.

## References

Adak A, Kang M, Anderson SL, Murray SC, Jarquin D, Wong RK, Katzfuß M (2023) Phenomic data-driven biological prediction of maize through field-based high-throughput phenotyping integration with genomic data. Journal of Experimental Botany 74:5307–5326

Adunola P, Tavares Flores E, Azevedo C, Casorzo G, Ghimire L, Ferrão LFV, Munoz PR (2024a) Phenomic-assisted selection: Assessment of the potential of near-infrared spectroscopy for blueberry breeding. The Plant Phenome Journal 7:e70010

Akdemir D, Isidro-Sánchez J (2019) Design of training populations for selective phenotyping in genomic prediction. Scientific Reports 9:1446

Alves-Carvalho S, Aubert G, Carrère S, Cruaud C, Brochot A-L, Jacquin F, Klein A, Martin C, Boucherot K, Kreplak J, da Silva C, Moreau S, Gamas P, Wincker P, Gouzy J, Burstin J (2015) Full-length de novo assembly of RNA-seq data in pea (Pisum sativum L.) provides a gene expression atlas and gives insights into root nodulation in this species. The Plant Journal 84:1–19

Annicchiarico P, Nazzicari N, Pecetti L, Romani M, Russi L (2019) Pea genomic selection for Italian environments. BMC genomics 20:603

Atanda SA, Steffes J, Lan Y, Al Bari MA, Kim JH, Morales M, Johnson JP, Saludares R, Worral H, Piche L, Ross A, Grusak M, Coyne C, McGee R, Rao J, Bandillo N (2022) Multi-trait genomic prediction improves selection accuracy for enhancing seed mineral concentrations in pea. Plant Genome 15:e20260

Aznar-Fernández T, Barilli E, Cobos MJ, Kilian A, Carling J, Rubiales D (2020) Identification of quantitative trait loci (QTL) controlling resistance to pea weevil (Bruchus pisorum) in a high-density integrated DArTseq SNP-based genetic map of pea. Sci Rep 10:33

Bari MAA, Zheng P, Viera I, Worral H, Szwiec S, Ma Y, Main D, Coyne CJ, McGee RJ, Bandillo N (2021) Harnessing Genetic Diversity in the USDA Pea Germplasm Collection Through Genomic Prediction. Frontiers in Genetics 12

Barilli E, Carrillo-Perdomo E, Cobos MJ, Kilian A, Carling J, Rubiales D (2020) Identification of potential candidate genes controlling pea aphid tolerance in a Pisum fulvum high-density integrated DArTseq SNP-based genetic map. Pest management science 76:1731–1742

Beji S, Fontaine V, Devaux R, Thomas M, Negro SS, Bahrman N, Siol M, Aubert G, Burstin J, Hilbert JL, Delbreil B, Lejeune-Hénaut I (2020) Genome-wide association study identifies favorable SNP alleles and candidate genes for frost tolerance in pea. BMC genomics 21:536

Bernardo R, Yu J (2007) Prospects for Genomewide Selection for Quantitative Traits in Maize. Crop Science 47:1082–1090

Bienvenu C, Garin V, Salas N, Théra K, Tekete ML, Sarathjith MC, Diallo C, Berger A, Calatayud C, De Bellis F (2025) Factors Influencing Phenomic Prediction: A Case Study on a Large Sorghum BCNAM Population. bioRxiv:2025.2002. 2004.636400

Bourgeois M, Jacquin F, Cassecuelle F, Savois V, Belghazi M, Aubert G, Quillien L, Huart M, Marget P, Burstin J (2011) A PQL (protein quantity loci) analysis of mature pea seed proteins identifies loci determining seed protein composition. PROTEOMICS 11:1581–1594

Brault C, Lazerges J, Doligez A, Thomas M, Ecarnot M, Roumet P, Bertrand Y, Berger G, Pons T, François P (2022) Interest of phenomic prediction as an alternative to genomic prediction in grapevine. Plant Methods 18:108

Browning BL, Browning SR (2007) Efficient multilocus association testing for whole genome association studies using localized haplotype clustering. Genetic Epidemiology 31:365–375

Burstin J, Gallardo K, Mir R, Varshney R, Duc G (2011) Improving protein content and nutrition quality. CABI:314–328

Burstin J, Salloignon P, Chabert-Martinello M, Magnin-Robert J-B, Siol M, Jacquin F, Chauveau A, Pont C, Aubert G, Delaitre C, Truntzer C, Duc G (2015) Genetic diversity and trait genomic prediction in a pea diversity panel. BMC genomics 16:105

Carpenter MA, Goulden DS, Woods CJ, Thomson SJ, Kenel F, Frew TJ, Cooper RD, Timmerman-Vaughan GM (2018) Genomic Selection for Ascochyta Blight Resistance in Pea. Front Plant Sci 9:1878

Champ M, Magrini M-B, Simon N, Guillou C, Schneider A, Huyghe C, Coord (2015) Les légumineuses pour des systèmes agricoles et alimentaires durables

Coyne CJ, Porter LD, Boutet G, Ma Y, McGee RJ, Lesné A, Baranger A, Pilet-Nayel M-L (2019) Confirmation of Fusarium root rot resistance QTL Fsp-Ps 2.1 of pea under controlled conditions. BMC Plant Biology 19:98

Crossa J, Jarquín D, Franco J, Pérez-Rodríguez P, Burgueño J, Saint-Pierre C, Vikram P, Sansaloni C, Petroli C, Akdemir D (2016) Genomic prediction of gene bank wheat landraces. G3: Genes, Genomes, Genetics 6:1819–1834

Crossa J, Pérez-Rodríguez P, Cuevas J, Montesinos-López O, Jarquín D, de los Campos G, Burgueño J, González-Camacho JM, Pérez-Elizalde S, Beyene Y, Dreisigacker S, Singh R, Zhang X, Gowda M, Roorkiwal M, Rutkoski J, Varshney RK (2017) Genomic Selection in Plant Breeding: Methods, Models, and Perspectives. Trends in Plant Science 22:961–975

Crosta M, Romani M, Nazzicari N, Ferrari B, Annicchiarico P (2023) Genomic prediction and allele mining of agronomic and morphological traits in pea (Pisum sativum) germplasm collections. Frontiers in Plant Science 14

Crosta M, Nazzicari N, Pecetti L, Notario T, Romani M, Ferrari B, Cabassi G, Annicchiarico P (2025) Genomic Selection for Pea Grain Yield and Protein Content in Italian Environments for Target and Non- Target Genetic Bases. International Journal of Molecular Sciences 26:2991

Dallinger HG, Löschenberger F, Bistrich H, Ametz C, Hetzendorfer H, Morales L, Michel S, Buerstmayr H (2023) Predictor bias in genomic and phenomic selection. Theoretical and Applied Genetics 136:235

DeSalvio AJ, Adak A, Murray SC, Jarquín D, Winans ND, Crozier D, Rooney WL (2024) Near-infrared reflectance spectroscopy phenomic prediction can perform similarly to genomic prediction of maize agronomic traits across environments. The Plant Genome 17:e20454

Desgroux A, L’Anthoëne V, Roux-Duparque M, Rivière J-P, Aubert G, Tayeh N, Moussart A, Mangin P, Vetel P, Piriou C, McGee RJ, Coyne CJ, Burstin J, Baranger A, Manzanares-Dauleux M, Bourion V, Pilet-Nayel M-L (2016) Genome-wide association mapping of partial resistance to Aphanomyces euteiches in pea. BMC genomics 17:124

De Verdal H, Segura V, Pot D, Salas N, Garin V, Rakotoson T, Raboin L-M, VomBrocke K, Dusserre J, Castro Pacheco SA (2024) Performance of phenomic selection in rice: Effects of population size and genotype-environment interactions on predictive ability. PloS one 19:e0309502

Duarte J, Riviere N, Baranger A, Aubert G, Burstin J, Cornet L, Lavaud C, Lejeune-Henaut I, Martinant J-P, Pichon J-P, Pilet-Nayel M-L, Boutet G (2014) Transcriptome sequencing for high throughput SNP development and genetic mapping in Pea. BMC genomics 15:126

Ellis N, Hofer J, Sizer-Coverdale E, Lloyd D, Aubert G, Kreplak J, Burstin J, Cheema J, Bal M, Chen Y, Deng S, Wouters RHM, Steuernagel B, Chayut N, Domoney C (2023) Recombinant inbred lines derived from wide crosses in Pisum. Scientific Reports 13:20408

Erbe M, Gredler B, Seefried FR, Bapst B, Simianer H (2013) A function accounting for training set size and marker density to model the average accuracy of genomic prediction. PLoS One 8:e81046

Fugeray-Scarbel A, Lemarié S (2024) The amplified effect of market size on innovation: A comparative analysis of pea and wheat seed value chains in France. Agricultural Systems 219:104051

Galán RJ, Bernal-Vasquez A-M, Jebsen C, Piepho H-P, Thorwarth P, Steffan P, Gordillo A, Miedaner T (2020) Integration of genotypic, hyperspectral, and phenotypic data to improve biomass yield prediction in hybrid rye. Theoretical and Applied Genetics 133:3001–3015

Gali KK, Sackville A, Tafesse EG, Lachagari VBR, McPhee K, Hybl M, Mikić A, Smýkal P, McGee R, Burstin J, Domoney C, Ellis THN, Tar’an B, Warkentin TD (2019) Genome-Wide Association Mapping for Agronomic and Seed Quality Traits of Field Pea (Pisum sativum L.). Frontiers in Plant Science 10

Gali KK, Jha A, Tar’an B, Burstin J, Aubert G, Bing D, Arganosa G, Warkentin TD (2024) Identification of QTLs associated with seed protein concentration in two diverse recombinant inbred line populations of pea. Front Plant Sci 15:1359117

Garcia-Abadillo J, Adunola P, Aguilar FS, Trujillo-Montenegro JH, Riascos JJ, Persa R, Isidro y Sanchez J, Jarquín D (2024) Sparse testing designs for optimizing predictive ability in sugarcane populations. Frontiers in Plant Science Volume 15–2024

Gilmour AR, Thompson R, Cullis BR (1995) Average information REML: an efficient algorithm for variance parameter estimation in linear mixed models. Biometrics:1440–1450

Gondalia N, Vashi R, Barot V, Sharma F, Anishkumar PK, Chatterjee M, Karmakar N, Gupta P, Sarker A, Kumar S, Sarkar A (2022) Genomic Designing for Abiotic Stress Tolerance in Pea (Pisum Sativum L.). In: Kole C (ed) Genomic Designing for Abiotic Stress Resistant Pulse Crops. Springer International Publishing, Cham, pp 45-113

Graciano RP, Peixoto MA, Leach KA, Suzuki N, Gustin JL, Settles AM, Armstrong PR, Resende Jr MFR (2025) Integrating phenomic selection using single-kernel near-infrared spectroscopy and genomic selection for corn breeding improvement. Theoretical and Applied Genetics 138:60

Haile TA, Heidecker T, Wright D, Neupane S, Ramsay L, Vandenberg A, Bett KE (2020) Genomic selection for lentil breeding: Empirical evidence. The Plant Genome 13:e20002

Heslot N, Jannink J-L, Sorrells ME (2015) Perspectives for Genomic Selection Applications and Research in Plants. Crop Science 55:1–12

Heuer C (2016) cpgen: Parallel Genomic Evaluations. R package https://github.com/cheuerde/cpgen.

Huang S, Gali KK, Arganosa GC, Tar᾿an B, Bueckert RA, Warkentin TD (2023) Breeding indicators for high-yielding field pea under normal and heat stress environments. Canadian Journal of Plant Science 103:259–269

Iglesias-García R, Prats E, Fondevilla S, Šatović Z, Rubiales D (2015) Quantitative Trait Loci Associated to Drought Adaptation in Pea (Pisum sativum L.). Plant Molecular Biology Reporter 33

Isidro J, Jannink J-L, Akdemir D, Poland J, Heslot N, Sorrells M (2015) Training set optimization under population structure in genomic selection. Theoretical and Applied Genetics 128:145–158

Jain S, Weeden NF, Kumar A, Chittem K, McPhee K (2015) Functional Codominant Marker for Selecting the Fw Gene Conferring Resistance to Fusarium Wilt Race 1 in Pea. Crop Science 55:2639–2646

Jarquín D, Crossa J, Lacaze X, Du Cheyron P, Daucourt J, Lorgeou J, Piraux F, Guerreiro L, Pérez P, Calus M, Burgueño J, de los Campos G (2014) A reaction norm model for genomic selection using high- dimensional genomic and environmental data. Theoretical and Applied Genetics 127:595–607

Jarquín D, Specht J, Lorenz A (2016) Prospects of genomic prediction in the USDA soybean germplasm collection: historical data creates robust models for enhancing selection of accessions. G3: Genes, Genomes, Genetics 6:2329–2341

Jha AB, Gali KK, Tar’an B, Warkentin TD (2017) Fine Mapping of QTLs for Ascochyta Blight Resistance in Pea Using Heterogeneous Inbred Families. Frontiers in Plant Science 8

Klein A, Houtin H, Rond-Coissieux C, Naudet-Huart M, Touratier M, Marget P, Burstin J (2020) Meta- analysis of QTL reveals the genetic control of yield-related traits and seed protein content in pea. Scientific Reports 10:15925

Klein A, Houtin H, Rond C, Marget P, Jacquin F, Boucherot K, Huart M, Rivière N, Boutet G, Lejeune-Hénaut I, Burstin J (2014) QTL analysis of frost damage in pea suggests different mechanisms involved in frost tolerance. Theoretical and Applied Genetics 127:1319–1330

Kreplak J, Madoui M-A, Cápal P, Novák P, Labadie K, Aubert G, Bayer PE, Gali KK, Syme RA, Main D, Klein A, Bérard A, Vrbová I, Fournier C, d’Agata L, Belser C, Berrabah W, Toegelová H, Milec Z, Vrána J, Lee H, Kougbeadjo A, Térézol M, Huneau C, Turo CJ, Mohellibi N, Neumann P, Falque M, Gallardo K, McGee R, Tar’an B, Bendahmane A, Aury J-M, Batley J, Le Paslier M-C, Ellis N, Warkentin TD, Coyne CJ, Salse J, Edwards D, Lichtenzveig J, Macas J, Doležel J, Wincker P, Burstin J (2019) A reference genome for pea provides insight into legume genome evolution. Nature Genetics 51:1411–1422

Kreplak J, Novak P, Robledillo LA, Aubert G, Imbert B, Kaur P, Gouil Q, Lopez-Roques C, Rodde N, Bouchez O, Tayeh N, Macas J, Burstin J (2025) A new genome assembly of the pea cultivar Cameor provides resources for functional genomics and genetics. bioRxiv:2025.2004.2001.645976

Laurençon M, Legrix J, Wagner M-H, Demilly D, Baron C, Rolland S, Ducournau S, Laperche A, Nesi N (2024) Genomic and phenomic predictions help capture low-effect alleles promoting seed germination in oilseed rape in addition to QTL analyses. Theoretical and Applied Genetics 137:156

Lavaud C, Lesné A, Leprévost T, Pilet-Nayel M-L (2024) Fine mapping of Ae-Ps4.5, a major locus for resistance to pathotype III of Aphanomyces euteiches in pea. Theoretical and Applied Genetics 137:47

Lecomte C, Richer V, Gauffreteau A, Jeuffroy M-H, Bouviala M, Brun C, Buridan C, Klein A, Lantoine F-X, Marchand D, Martin J, Naudet-Huart M, Tayeh N, Touratier M, Valdrini J-M, Walczak P, Burstin J (2023) Combining a multi-environment trial and a diagnosis method to assess potential yield and main limiting factors of three highly different pea types. European Journal of Agronomy 146:126823

Lejeune-Henaut I, Hanocq E, Bethencourt L, Fontaine V, Delbreil B, Morin J, Petit A, Devaux R, Boilleau M, Stempniak JJ, Thomas M, Laine AL, Foucher F, Baranger A, Burstin J, Rameau C, Giauffret C (2008) The flowering locus Hr colocalizes with a major QTL affecting winter frost tolerance in Pisum sativum L. Theoretical and Applied Genetics 116:1105–1116

Leprévost T, Boutet G, Lesné A, Rivière J-P, Vetel P, Glory I, Miteul H, Le Rat A, Dufour P, Regnault-Kraut C, Sugio A, Lavaud C, Pilet-Nayel M-L (2023) Advanced backcross QTL analysis and comparative mapping with RIL QTL studies and GWAS provide an overview of QTL and marker haplotype diversity for resistance to Aphanomyces root rot in pea (Pisum sativum). Frontiers in Plant Science 14

Li Y, Kaur S, Pembleton LW, Valipour-Kahrood H, Rosewarne GM, Daetwyler HD (2022) Strategies of preserving genetic diversity while maximizing genetic response from implementing genomic selection in pulse breeding programs. TAG Theoretical and applied genetics Theoretische und angewandte Genetik 135:1813–1828

Ligges U, Short T, Kienzle P, Schnackenberg S, Billinghurst D, Borchers H-W, Carezia A, Dupuis P, Eaton JW, Farhi E (2015) Package ‘signal’. R Foundation for Statistical Computing

Liu N, Lyu X, Zhang X, Zhang G, Zhang Z, Guan X, Chen X, Yang X, Feng Z, Gao Q, Shi W, Deng Y, Sheng K, Ou J, Zhu Y, Wang B, Bu Y, Zhang M, Zhang L, Zhao T, Gong Y (2024) Reference genome sequence and population genomic analysis of peas provide insights into the genetic basis of Mendelian and other agronomic traits. Nature Genetics 56:1964–1974

Magrini M-B, Anton M, Cholez C, Corre-Hellou G, Duc G, Jeuffroy M-H, Meynard J-M, Pelzer E, Voisin A-S, Walrand S (2016) Why are grain-legumes rarely present in cropping systems despite their environmental and nutritional benefits? Analyzing lock-in in the French agrifood system. Ecological Economics 126:152–162

McGaugh SE, Lorenz AJ, Flagel LE (2021) The utility of genomic prediction models in evolutionary genetics. Proceedings Biological sciences 288:20210693

Meuwissen TH, Hayes BJ, Goddard ME (2001) Prediction of total genetic value using genome-wide dense marker maps. Genetics 157:1819–1829

Meyenberg C, Braun V, Longin CFH, Thorwarth P (2024) Feature engineering and parameter tuning: improving phenomic prediction ability in multi-environmental durum wheat breeding trials. Theoretical and Applied Genetics 137:188

Michel S, Löschenberger F, Ametz C, Pachler B, Sparry E, Bürstmayr H (2019) Combining grain yield, protein content and protein quality by multi-trait genomic selection in bread wheat. Theoretical and Applied Genetics 132:2767–2780

Osuna-Caballero S, Rubiales D, Annicchiarico P, Nazzicari N, Rispail N (2024) Genomic prediction for rust resistance in pea. Frontiers in Plant Science 15

Paleari L, Tondelli A, Cattivelli L, Igartua E, Casas AM, Visioni A, Schulman AH, Rossini L, Waugh R, Russell J, Confalonieri R (2025) Extending genomic prediction to future climates through crop modelling. A case study on heading time in barley. Agricultural and Forest Meteorology 368:110560

Parihar AK, Kumar J, Gupta DS, Lamichaney A, Naik SJ S, Singh AK, Dixit GP, Gupta S, Toklu F (2022) Genomics Enabled Breeding Strategies for Major Biotic Stresses in Pea (Pisum sativum L.). Frontiers in Plant Science 13

R Core team (2021) R: A language and environment for statistical computing. R Foundation for Statistical Computing, Vienna, Austria. https://www.R-project.org/.

Rincent R, Charpentier JP, Faivre-Rampant P, Paux E, Le Gouis J, Bastien C, Segura V (2018) Phenomic Selection Is a Low-Cost and High-Throughput Method Based on Indirect Predictions: Proof of Concept on Wheat and Poplar. G3 (Bethesda, Md) 8:3961–3972

Rincent R, Solin J, Lorenzi A, Nune s L, Griveau Y, Pirus L, Kermarrec D, Bauland C, Reymond M, Moreau L (2025) Using phenomic selection to predict hybrid values with NIR spectra measured on the parental lines: proof of concept on maize. Theoretical and Applied Genetics 138:28

Robert P, Auzanneau J, Goudemand E, Oury FX, Rolland B, Heumez E, Bouchet S, Le Gouis J, Rincent R (2022a) Phenomic selection in wheat breeding: identification and optimisation of factors influencing prediction accuracy and comparison to genomic selection. TAG Theoretical and applied genetics Theoretische und angewandte Genetik 135:895–914

Robert P, Goudemand E, Auzanneau J, Oury FX, Rolland B, Heumez E, Bouchet S, Caillebotte A, Mary-Huard T, Le Gouis J, Rincent R (2022b) Phenomic selection in wheat breeding: prediction of the genotype-by-environment interaction in multi-environment breeding trials. TAG Theoretical and applied genetics Theoretische und angewandte Genetik 135:3337–3356

Roscher-Ehrig L, Weber SE, Abbadi A, Malenica M, Abel S, Hemker R, Snowdon RJ, Wittkop B, Stahl A (2024) Phenomic selection for hybrid rapeseed breeding. Plant Phenomics 6:0215

Saludares RA, Atanda SA, Piche L, Worral H, Dariva F, McPhee K, Bandillo N (2024) Multi-trait multi- environment genomic prediction of preliminary yield trial in pulse crop. The Plant Genome 17:e20496

Sandhu KS, Mihalyov PD, Lewien MJ, Pumphrey MO, Carter AH (2021) Combining genomic and phenomic information for predicting grain protein content and grain yield in spring wheat. Frontiers in plant science 12:613300

Schaid DJ (2010) Genomic similarity and kernel methods I: advancements by building on mathematical and statistical foundations. Human heredity 70:109–131

Singh H, Asija S, Sharma K, Koul B, Tiwari S (2023) Genetic Improvement of Pea (Pisum sativum L.) for Food and Nutritional Security. In: Tiwari S, Koul B (eds) Genetic Engineering of Crop Plants for Food and Health Security: Volume 1. Springer Nature Singapore, Singapore, pp 1–37

Smits G, Jordaan EM (2002) Improved SVM regression using mixtures of kernels

Smýkal P, K. Varshney R, K. Singh V, Coyne CJ, Domoney C, Kejnovský E, Warkentin T (2016) From Mendel’s discovery on pea to today’s plant genetics and breeding. Theoretical and Applied Genetics 129:2267–2280

Stevens A, Ramirez–Lopez L (2013) An introduction to the prospectr package

Tafesse EG, Gali KK, Lachagari VBR, Bueckert R, Warkentin TD (2021) Genome-Wide Association Mapping for Heat and Drought Adaptive Traits in Pea. Genes 12:1897

Tar’an B, Warkentin T, Somers DJ, Miranda D, Vandenberg A, Blade S, Bing D (2004) Identification of quantitative trait loci for grain yield, seed protein concentration and maturity in field pea (Pisum sativum L.). Euphytica 136:297–306

Tayeh N, Aluome C, Falque M, Jacquin F, Klein A, Chauveau A, Bérard A, Houtin H, Rond C, Kreplak J, Boucherot K, Martin C, Baranger A, Pilet-Nayel M-L, Warkentin TD, Brunel D, Marget P, Le Paslier M-C, Aubert G, Burstin J (2015a) Development of two major resources for pea genomics: the GenoPea 13.2K SNP Array and a high-density, high-resolution consensus genetic map. The Plant Journal 84:1257–1273

Tayeh N, Aubert G, Pilet-Nayel M-L, Lejeune-Hénaut I, Warkentin TD, Burstin J (2015b) Genomic Tools in Pea Breeding Programs: Status and Perspectives. Frontiers in Plant Science 6

Tayeh N, Klein A, Le Paslier M-C, Jacquin F, Houtin H, Rond C, Chabert-Martinello M, Magnin-Robert J-B, Marget P, Aubert G, Burstin J (2015c) Genomic Prediction in Pea: Effect of Marker Density and Training Population Size and Composition on Prediction Accuracy. Frontiers in Plant Science 6

Teressa T, Semahegn Z, Bejiga T (2021) Multi environments and genetic-environmental interaction (GxE) in plant breeding and its challenges: a review article. International Journal of Research Studies in Agricultural Sciences 7:11–18

VanRaden PM (2008) Efficient Methods to Compute Genomic Predictions. Journal of Dairy Science 91:4414–4423

Wang F, Feldmann MJ, Runcie DE (2025) Why Accuracy Metrics Fall Short in Comparing Phenomic and Genomic Prediction Models. bioRxiv:2025.2001. 2009.632209

Wang K, Abid MA, Rasheed A, Crossa J, Hearne S, Li H (2023) DNNGP, a deep neural network-based method for genomic prediction using multi-omics data in plants. Molecular Plant 16:279–293

Xu Y, Liu X, Fu J, Wang H, Wang J, Huang C, Prasanna BM, Olsen MS, Wang G, Zhang A (2020) Enhancing Genetic Gain through Genomic Selection: From Livestock to Plants. Plant Communications 1:100005

Yang T, Liu R, Luo Y, Hu S, Wang D, Wang C, Pandey MK, Ge S, Xu Q, Li N (2022) Improved pea reference genome and pan-genome highlight genomic features and evolutionary characteristics. Nature genetics 54:1553–1563

Zhu X, Leiser WL, Hahn V, Würschum T (2021) Phenomic selection is competitive with genomic selection for breeding of complex traits. The Plant Phenome Journal 4:e20027

