## Supplementary material for "Combining phenomic and genomic selection for pea breeding improvement": e.g. Supplemental figures

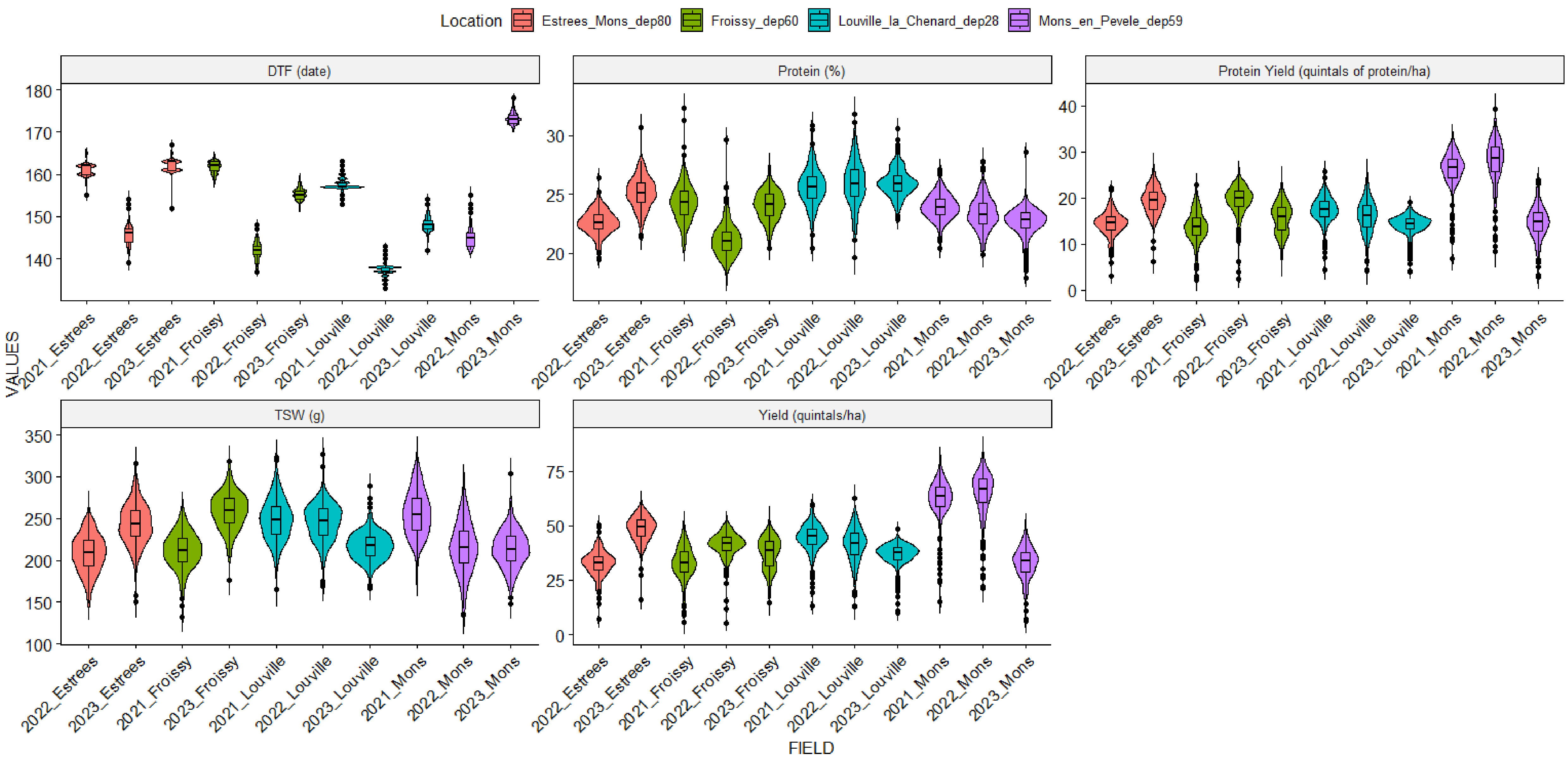


**Fig.S1** Distribution of phenotypes by environment, as observed in field trials at four locations from 2021 to 2023. *DTF* days to flowering (date), *TSW* thousand seed weight (grams), *Yield* seed yield at maturity (quintals per hectare), *Protein* seed protein content at maturity (percentage of dry matter), *Protein_Yield* seed protein yield at maturity (quintals of protein per hectare).

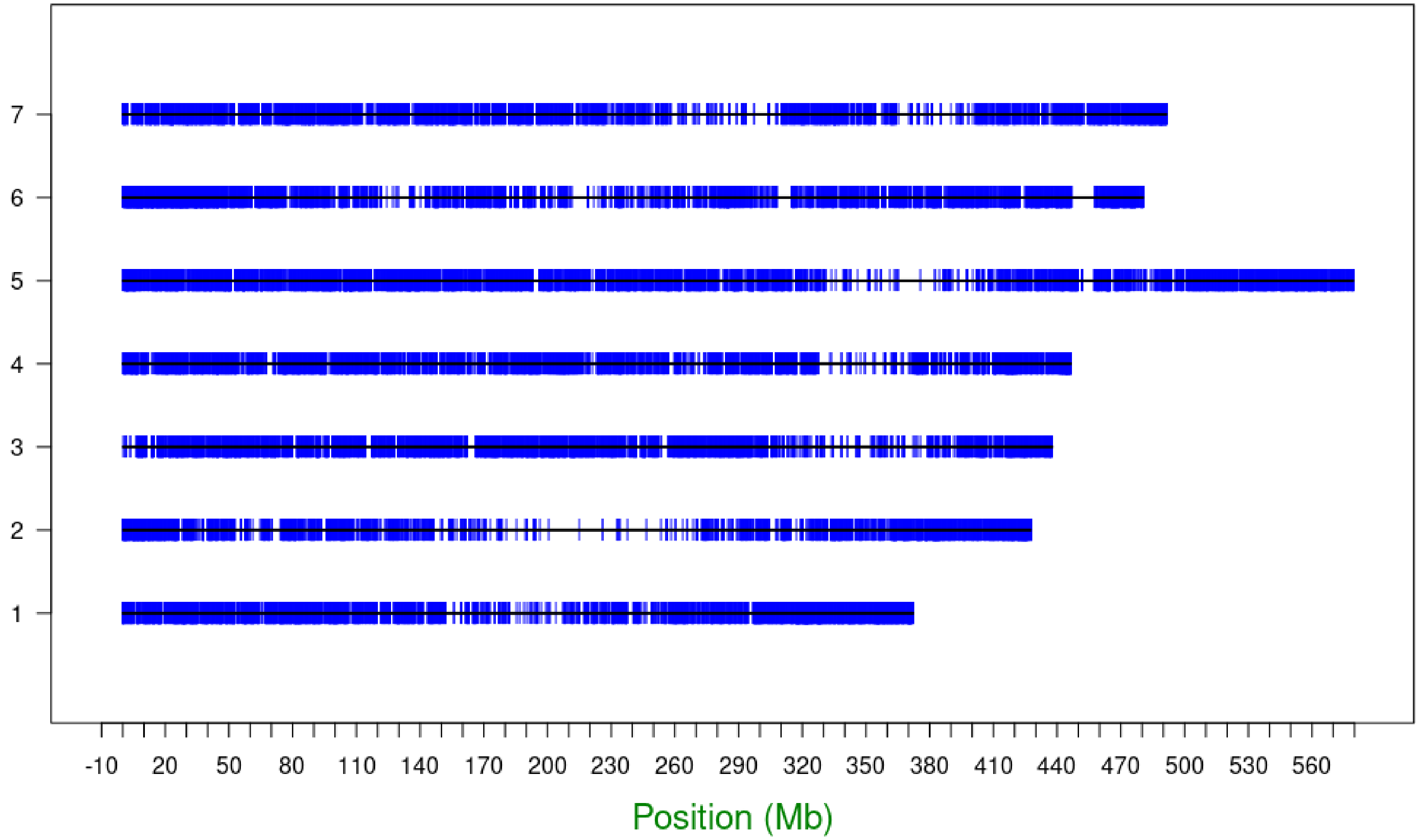


**Fig.S2** Distribution of SNPs along each chromosome of Caméor V1. Each blue line represents the position of a SNP marker in Mb on a chromosome

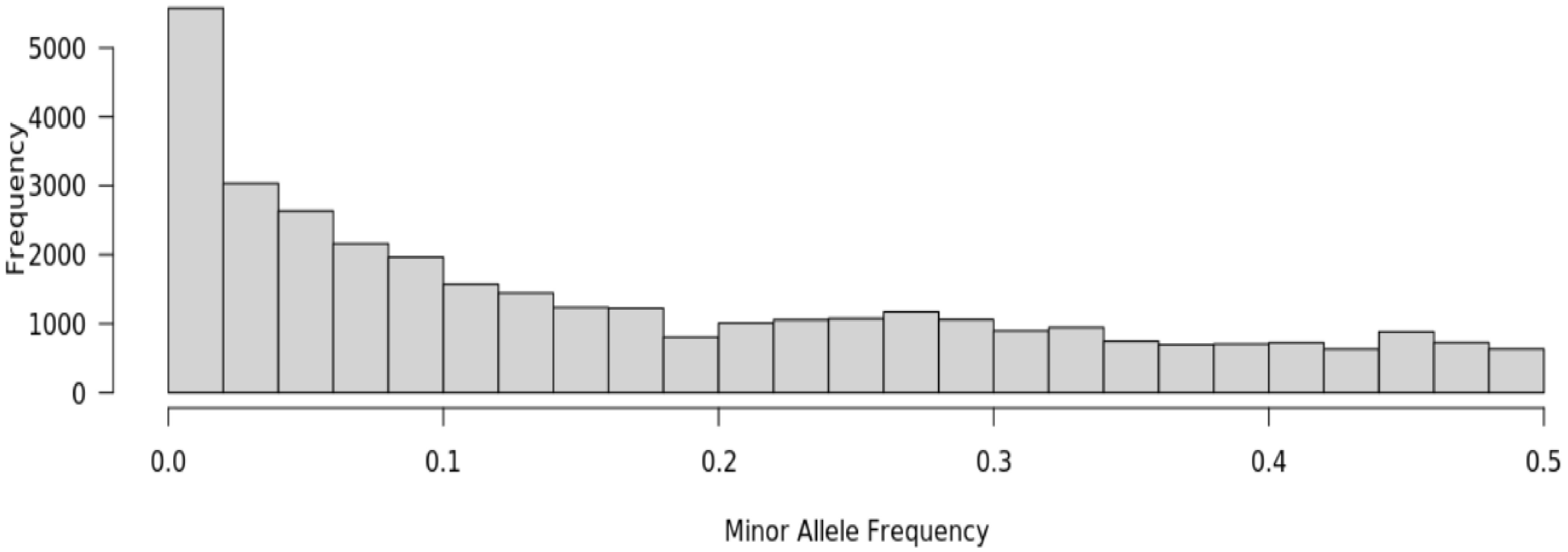


**Fig.S3** Minority allele frequency (MAF) distribution of SNPs markers

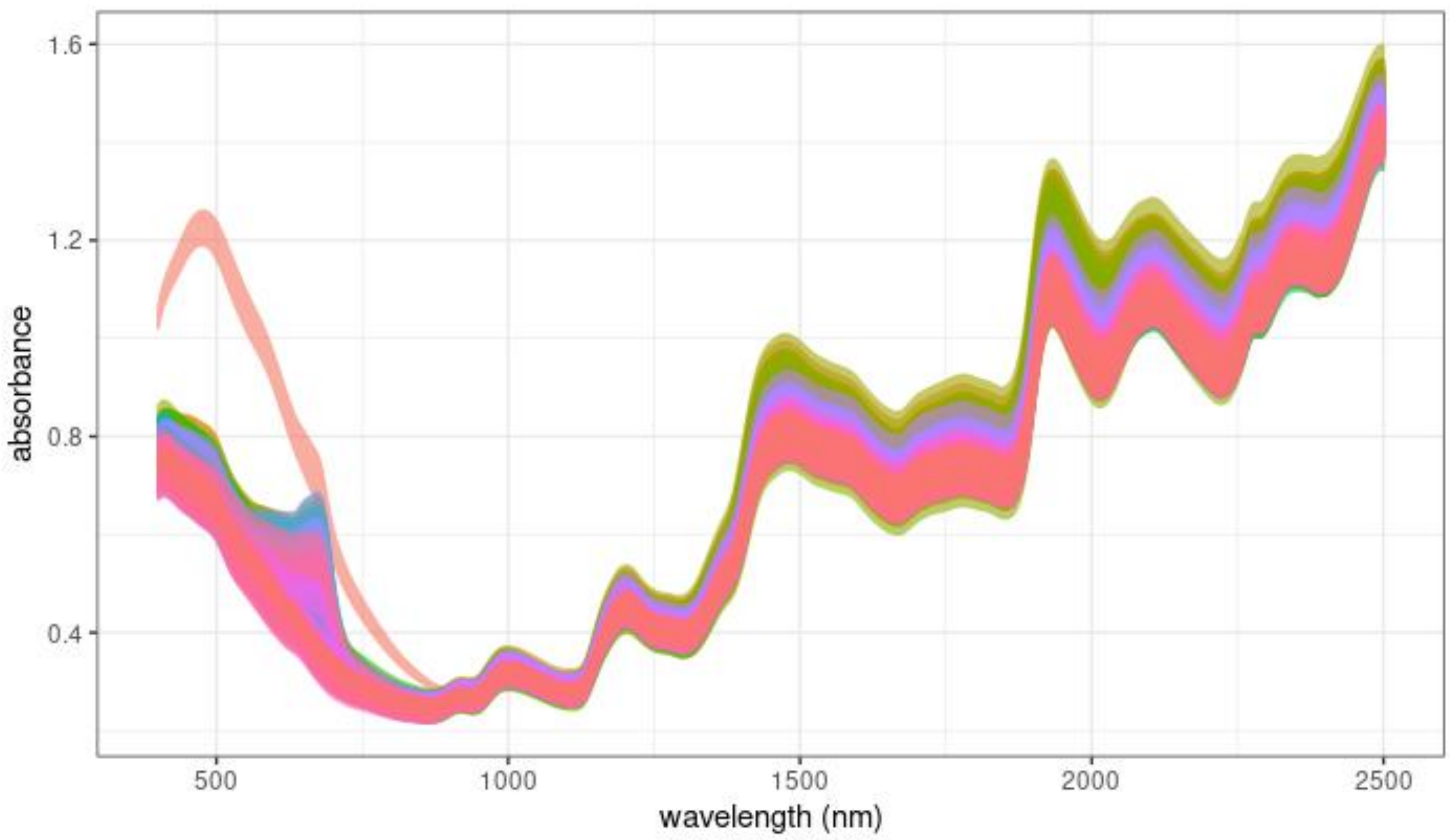


**Fig.S4** NIRS absorbance spectra from the seeds of the 288-acession panel, collected in all environments. Each colored line represents the raw spectrum for a different pea line and its absorbance values (y-axis) across a range of wavelengths from 400 nm to 2,499.5 nm (x-axis).
